# Virtual-cell models compress unseen intervention geometry through a target-specific generalization bottleneck

**DOI:** 10.64898/2026.08.21.746243

**Authors:** Yongqi Huang, Hanzhi Wang, Parker Wilson

## Abstract

Predictive models of cellular perturbation are often judged by how closely they reconstruct molecular states after unseen interventions. We show that high state-level similarity can coexist with loss of the relationships that distinguish perturbations, a failure we term Intervention Geometry Compression (IGC). Across established models and perturbation settings, unseen interventions show weakened global and local geometry, reduced between-intervention variance and spectral collapse. The failure is not primarily explained by response-space capacity. Instead, diagnostic projections localize much of the missing geometry to a small number of residual response directions learned from seen interventions; these directions outperform complexity-matched random subspaces and replicate in an independent Jiang perturbation resource. Polarity captures part, but not all, of this continuous orientation signal. Time-resolved analyses further show that correct trajectory entry markedly improves downstream propagation, while a held target’s own early empirical response rapidly reveals endpoint orientation. Finally, same-target empirical anchoring transfers intervention identity across contexts far more effectively than increasing exposure to other interventions. These results identify intervention-coordinate assignment as an information bottleneck in virtual-cell generalization and support a design principle: empirically anchor intervention identity, then use models to generalize anchored effects across cellular contexts.

## Introduction

Large-scale single-cell perturbation profiling has created the possibility of learning predictive models that map genetic interventions to cellular molecular states. In the most ambitious formulation, a virtual-cell model should predict the response to an intervention that was never observed during training^1^, allowing experimental programs to prioritize targets before measuring them directly. Large-scale benchmarking has highlighted that performance depends strongly on both the generalization axis and evaluation metric^2^, while perturbation-specific and foundation models increasingly target unseen genetic interventions and are commonly evaluated using reconstruction measures such as expression correlation^3–6^. At the same time, stronger baselines and stricter evaluation protocols can substantially alter model rankings^7^, motivating richer representations of intervention identity, including graph- and language-derived gene embeddings^8^.

Yet the central use case of perturbation prediction is not simply to generate a transcriptome that looks perturbed^9^. Models are expected to distinguish candidate interventions, preserve mechanistic similarities and differences^10^, rank targets^11^ and eventually guide experimental decisions^12^. These applications depend on intervention identity: perturbations that produce distinct biological responses must remain distinct in the predicted response space. A model can therefore achieve a high state-level correlation while failing at the more consequential task of determining which intervention produced which response^9^.

This distinction follows directly from the structure of perturbational expression data. A perturbed state contains a large control-state component, responses shared across many interventions and a smaller intervention-specific component^9,13^. Conventional state-level similarity can consequently remain high even when intervention-specific effects are poorly recovered. Related benchmarking studies have similarly shown that simple mean-response baselines can rival or exceed foundation-model predictions under commonly used perturbation metrics^14^. We therefore asked whether current perturbation models preserve the relative organization of interventions when they generalize to genuinely unseen targets. Recent benchmarking efforts have begun to emphasize perturbation ordering and rank-based evaluation beyond pointwise reconstruction error^15^. Here, we extend this perspective to the full pairwise geometry, local neighborhoods, variance structure and spectral dimensionality of unseen intervention responses.

Across multiple perturbation settings, we identify a recurrent failure mode that we term Intervention Geometry Compression (IGC). Models that reconstruct plausible absolute states map unseen interventions into a response landscape with weakened pairwise relationships, degraded local neighborhoods, contracted between-intervention variance and reduced spectral dimensionality. Seen-versus-unseen analyses, response-space reconstruction and architecture controls point to incorrect assignment of unseen interventions to response directions, rather than insufficient output capacity alone, as the dominant explanation.

This localization raised a more specific question: is the mapping error distributed across the full expression space or concentrated in a compact set of coordinates? Residual directions learned exclusively from seen interventions showed that one or two held-target coordinates recover a large fraction of missing geometry in K562, and the same low-dimensional pattern reproduced across IFNB, IFNG and INS perturbations in the Jiang pathway Perturb-seq resource^23^. Complexity-matched random-subspace controls show that the result is not a generic consequence of granting diagnostic access to a few dimensions. Polarity contributes useful information, but continuous coordinates provide substantially more, motivating a low-dimensional continuous intervention-orientation code rather than a literal two-bit representation.

Time-resolved perturbation data^16^ provided a dynamic test of what these intervention-specific coordinates do and when they become observable. True intermediate perturbation states and correct trajectory entry strongly improve later intervention geometry, indicating that downstream propagation capacity can be present even when an unseen intervention enters the wrong trajectory. Endpoint orientation axes were then fixed using training sources only, allowing us to ask how rapidly a held target’s own early perturbational response reveals its later orientation. Because this assay observes the held target after perturbation, it is an empirical observability diagnostic rather than a zero-shot predictor.

Recent work has shown that an empirical response of the same intervention can be reused to predict its effect in a missing recipient context^17^. The information value of such target-specific anchoring, however, has not been directly compared with simply observing more other interventions. A factorial experiment reveals a strong asymmetry: increasing non-target intervention coverage from 10% to 90% provides essentially no geometric rescue for a fixed unseen target set. An empirical response of the same target in another context provides a large, identity-specific transferable anchor. The advantage replicates in both K562-to-RPE1 and RPE1-to-K562^18^ transfer and independently in the Frangieh multi-context Perturb-CITE-seq system^20^. The resulting division of labour is simple: empirically anchor intervention identity where possible, and allocate model generalization primarily to the broader space of cellular contexts.

## Results

### Conventional perturbation metrics obscure intervention-specific prediction failure

To determine whether conventional reconstruction metrics capture intervention identity, we first separated perturbation prediction into progressively more intervention-specific quantities. Conceptually, a perturbed state can be decomposed into the control-state background, a response shared across interventions and an intervention-specific component^9^ (Fig. 1a,b). If the first two components dominate expression variance, a model can achieve high absolute-state similarity without recovering the component that distinguishes one perturbation from another.

**Figure 1.**
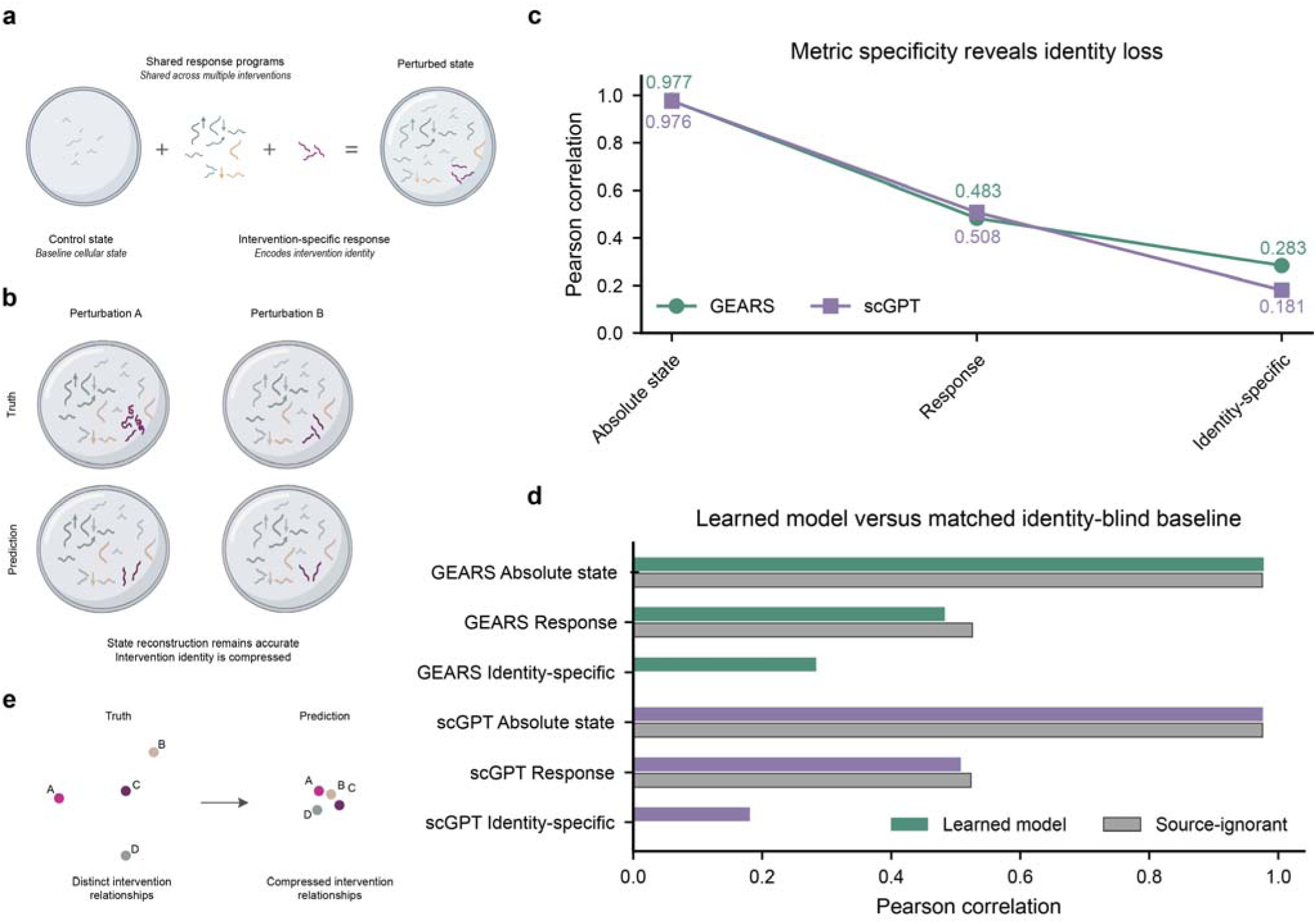
Conventional perturbation metrics obscure intervention-specific prediction failure. a, Conceptual decomposition of a perturbed state into control-state background, a response shared across interventions and an intervention-specific component. b, Schematic showing how overall state reconstruction can remain accurate even when the intervention-specific component is compressed. c, Fold-mean Pearson correlations for GEARS and scGPT evaluated at the absolute perturbed state, control-relative response and reference-mean-centered intervention-specific residual. d, Learned-model performance compared with a matched source-ignorant baseline that assigns every intervention the reference-derived mean response; the baseline has zero intervention-specific residual by construction. e, Conceptual transition from independent state reconstruction to relational intervention-geometry evaluation.

GEARS^3^ provided a direct example of this metric hierarchy. Absolute perturbed-state Pearson correlation was 0.977, but performance fell to 0.483 when evaluation was restricted to the control-relative perturbation response and to 0.283 for the intervention-specific residual component. scGPT^4^ showed the same hierarchy: absolute-state correlation was 0.976, response correlation was 0.508 and intervention-specific residual correlation fell to 0.181 (Fig. 1c). Thus, both established models remained highly accurate at the full-state level while losing substantially more of the component that distinguishes interventions.

A stronger demonstration came from a source-ignorant predictor that assigned all interventions the same reference-derived mean perturbation response. In the GEARS evaluation, this identity-blind baseline reached an absolute-state Pearson correlation of 0.975, nearly matching GEARS at 0.977, and its control-relative response correlation of 0.526 exceeded the learned model’s 0.483; its intervention-specific residual correlation was zero by construction. The same pattern reproduced for scGPT: the source-ignorant baseline reached 0.976 for absolute state and 0.524 for response, compared with 0.976 and 0.508 for scGPT, while its residual correlation again remained zero (Fig. 1d). Thus, strong conventional reconstruction metrics can arise in the complete absence of intervention-specific information.

These observations motivated a relational evaluation. Rather than asking whether each predicted state independently resembles its observed counterpart, we asked whether the model preserves which interventions are similar, which are distinct and how their response directions are organized relative to one another (Fig. 1e). This intervention-geometry perspective forms the basis of the subsequent analyses. Detailed metric decomposition and source-ignorant controls are provided in Supplementary Fig. 1, with artifact-safe geometry controls in Supplementary Fig. 2.

### Virtual-cell predictions compress the geometry of unseen interventions

Across large-scale Perturb-seq settings^18,19^, observed responses occupied a broad and heterogeneous landscape. OOF predictions for unseen interventions instead concentrated into a smaller set of directions (Fig. 2a). This qualitative narrowing suggested that conventional state reconstruction could coexist with loss of the organization that distinguishes interventions.

**Figure 2.**
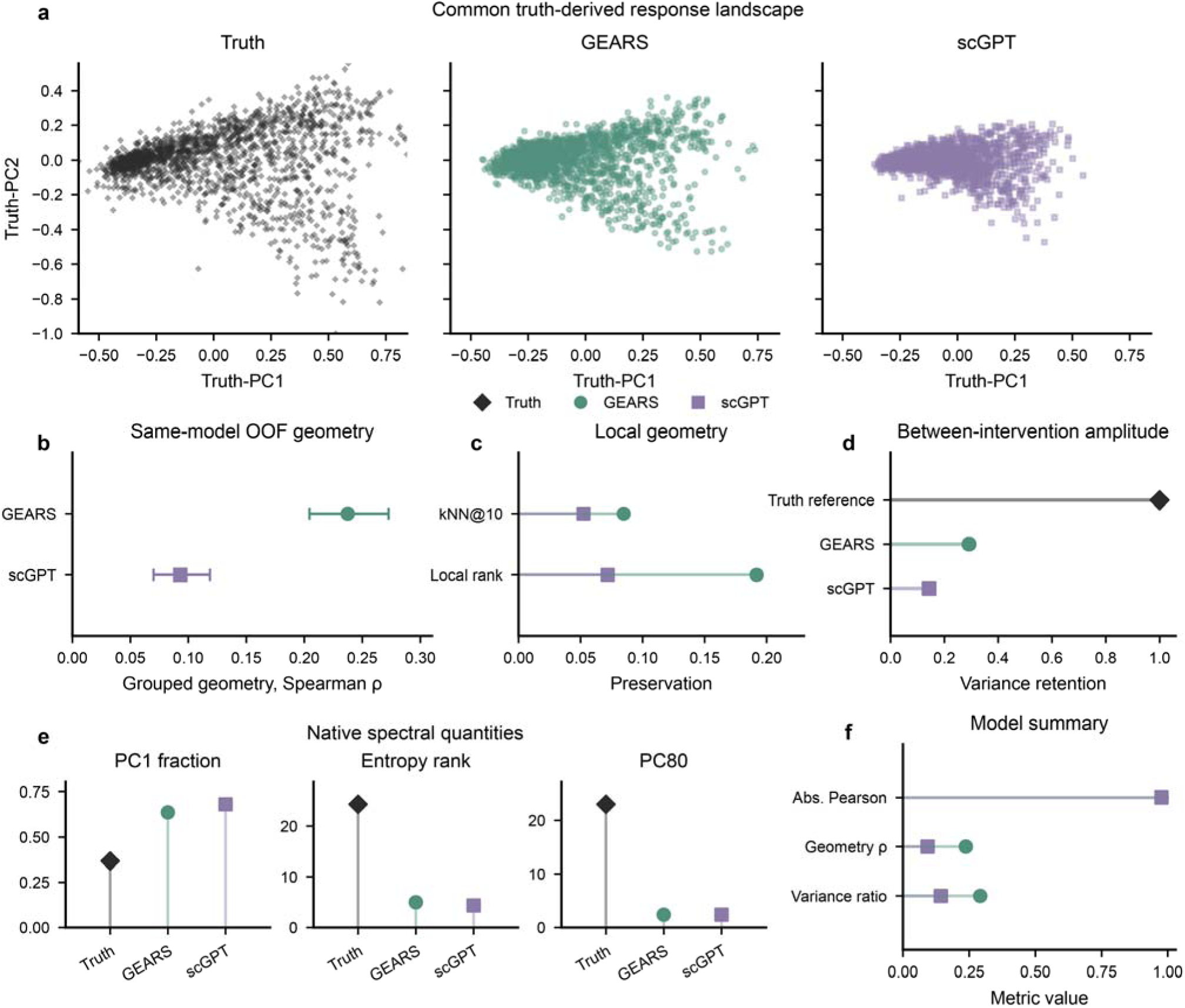
Virtual-cell predictions compress the geometry of unseen interventions. a, Observed and OOF-predicted intervention responses projected into a common truth-derived two-dimensional response basis with matched axes for Truth, GEARS and scGPT. b, Artifact-safe same-model grouped intervention geometry, defined as the Spearman correlation between predicted and observed pairwise cosine-distance ranks; error bars show source-bootstrap 95% confidence intervals. c, Local geometry measured by kNN@10 overlap and local distance-rank preservation. d, Between-intervention variance retention relative to the observed response reference (=1). e, Native spectral quantities: PC1 variance fraction, entropy effective rank and the number of components required to explain 80% of variance (PC80). f, Cross-model summary of absolute-state Pearson correlation, grouped intervention geometry and variance retention. Full matched established-model audits are shown in Supplementary Fig. 4.

To quantify this structure without introducing cross-model artifacts, geometry was computed only among predictions generated by the same held-out model and then aggregated across held-out groups. Under this artifact-safe evaluation, intervention geometry remained weak for both established models (Fig. 2b). GEARS retained measurable ordering among unseen interventions, with grouped geometry of 0.237 (95% bootstrap confidence interval, 0.204-0.273). scGPT retained substantially less geometry, at 0.093 (0.070-0.119). The phenotype is therefore a continuum rather than a binary success or failure, but it is reproduced across two established model classes.

Compression was also evident locally. GEARS achieved a k-nearest-neighbor overlap of 0.085 and a local distance-rank statistic of 0.192, compared with 0.052 and 0.072 for scGPT, respectively (Fig. 2c). Predictions retained only 29.2% of observed between-intervention response variance for GEARS and 14.3% for scGPT (Fig. 2d), showing that distinct perturbations were represented as too similar to one another.

The strongest signature arose from the response spectrum. In GEARS, the first predicted principal component captured 63.1% of between-intervention variation compared with 37.0% in matched observed responses; for scGPT, the corresponding values were 68.2% and 36.8% (Fig. 2e). Entropy-based effective rank fell from 23.89 to 5.07 for GEARS and from 24.33 to 4.35 for scGPT. The number of components required to explain 80% of variation contracted from 22.4 to 2.4 for GEARS and from 23.0 to 2.4 for scGPT. Thus, unseen responses are not merely smaller in magnitude: in both established models, a high-dimensional biological landscape is compressed into a few dominant predicted directions. The full matched GEARS and scGPT audit is provided in Supplementary Fig. 4.

A dedicated artifact audit was essential for interpreting these quantities. Naively concatenating predictions generated by different held-out models can manufacture apparent geometry even for a source-ignorant null predictor. In our audit, stitched evaluation produced a geometry of 1.0 for a null predictor. The correct same-model grouped calculation returned zero (Supplementary Fig. 2). All geometry values used in the main text therefore derive from same-model grouped evaluation. Bootstrap and pseudoreplicate reliability analyses further support that IGC cannot be attributed solely to measurement instability (Supplementary Fig. 3).

### Geometry compression reflects unseen-intervention mapping failure rather than limited response capacity

A simple lack of model capacity did not explain geometry compression. If a model were unable to represent the response landscape itself, geometry should be poor even for interventions encountered during training. Instead, a large Transformer preserved seen-intervention geometry in RPE1 cells (ρ = 0.741) but collapsed under source-disjoint evaluation of unseen interventions (ρ = 0.013), a gap of 0.729 (95% confidence interval, 0.706-0.749; Fig. 3a). The same architecture can therefore encode rich intervention structure while failing to transfer that structure to new intervention identities.

**Figure 3.**
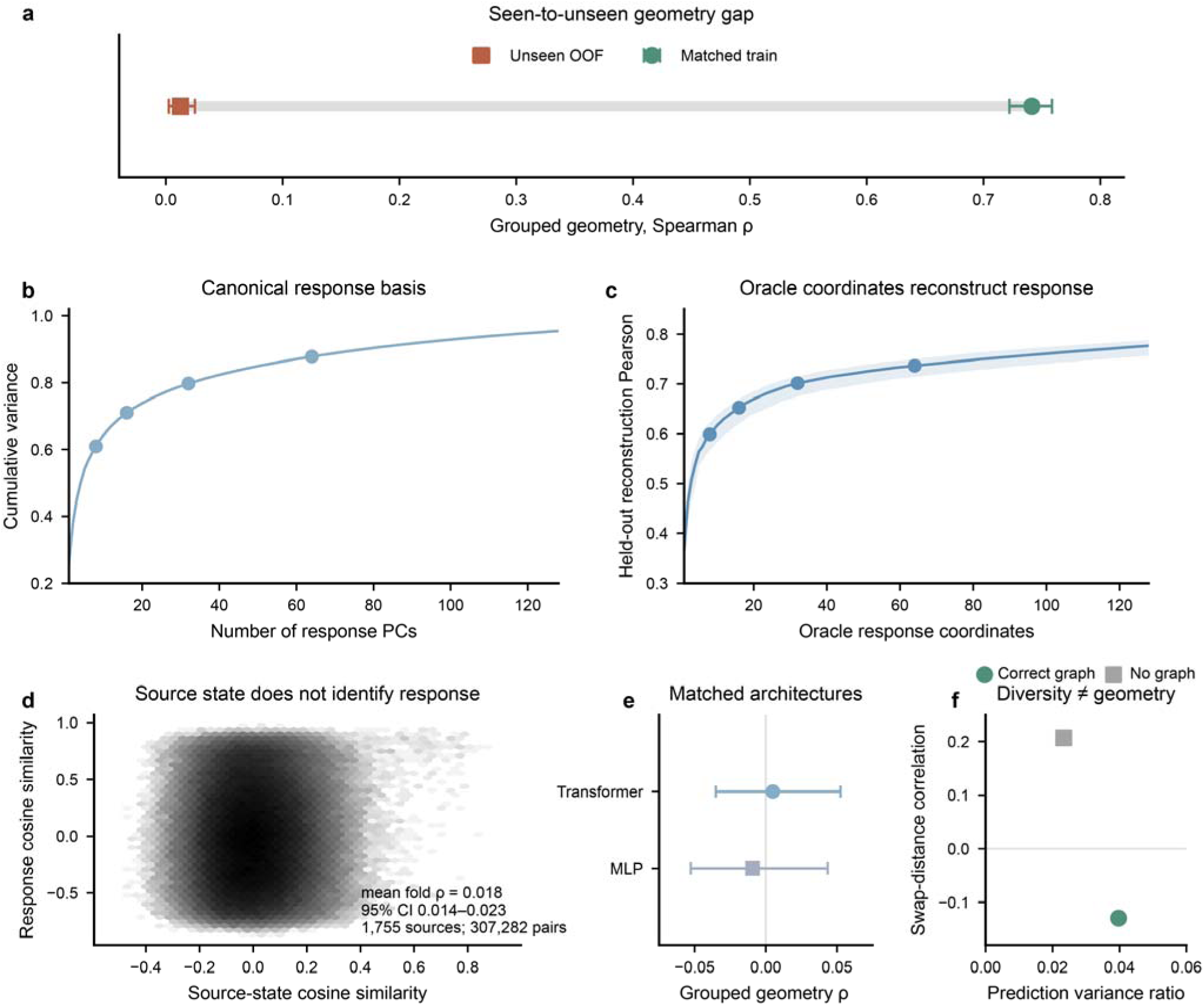
Geometry compression reflects unseen-intervention mapping failure rather than limited response capacity. a, Matched seen-intervention versus source-disjoint unseen-intervention geometry for a large Transformer in RPE1; error bars show 95% bootstrap confidence intervals. b, Cumulative variance captured by the canonical training-reference response PCA. c, Held-out response reconstruction when the correct oracle response-basis coordinates are supplied; the line shows the five-fold mean and shading the foldwise range. d, Density of 307,282 leakage-safe within-fold source pairs comparing EstablishedOBS71 control-state cosine similarity with true response cosine similarity; the annotation reports the canonical mean fold Spearman correlation and its 95% confidence interval. e, Matched Transformer-versus-MLP grouped-geometry control with 95% bootstrap confidence intervals. f, Anti-collapse control comparing prediction variance ratio with swap-distance correlation, illustrating that greater output diversity does not imply correct intervention geometry. Additional capacity and identifiability controls are shown in Supplementary Fig. 5.

The biological response space was also amenable to compact representation. Principal-component analysis of the training-side strict-trans response matrix showed that the first 8 components captured 60.9% of response variance and the first 16 components captured 71.0% (Fig. 3b). Perturbation responses therefore exhibited substantial low-dimensional structure, although a non-negligible fraction of variation remained outside the leading components.

Representability, however, does not establish predictability. Supplying the correct response-basis coordinates for held-out perturbations enabled accurate reconstruction of the corresponding responses (Fig. 3c). The response space itself is therefore representable; the harder step is assigning an unseen intervention to the correct location within that space.

Source-side information showed the opposite pattern. An established control-state representation had only negligible correspondence with the true intervention-specific response direction (Spearman ρ = 0.018, 95% confidence interval 0.014-0.023; Fig. 3d). Thus, the response space is representable, but the available intervention-side information provides almost no discrimination of where an unseen intervention should be placed within it.

We next tested a more direct possibility: whether spontaneous target-specific expression fluctuations across unperturbed single cells could act as natural perturbations and thereby provide the missing sender-side anchor. Across 11,485 RPE1 control cells, target–response co-fluctuation showed an apparent correspondence with the full control-relative perturbation response (mean cosine = 0.200), but this signal disappeared when evaluation was restricted to the intervention-specific residual (0.0057, 95% CI −0.0129 to 0.0230) and after residualizing major cell-state variation (−0.0018, 95% CI −0.0041 to 0.0004; Supplementary Fig. 5e,f). A fully source-disjoint calibration learned from reference targets likewise produced essentially zero held-out response correspondence and failed to restore intervention geometry (grouped ρ = 0.0073, 95% CI 0.0006–0.0143; Supplementary Fig. 5g,h). Thus, abundant observational heterogeneity captured shared cellular programs but did not provide a reliable target-specific sender anchor for unseen interventions.

Besides, the generalization failure was not specific to Transformer architecture. Under matched evaluation, replacing the Transformer with a multilayer perceptron did not support a Transformer-specific disadvantage (Fig. 3e and Supplementary Fig. 5). Finally, explicitly encouraging greater output diversity increased response dispersion without restoring the correct intervention relationships (Fig. 3f). Increasing diversity is therefore not sufficient to recover biological identity: different is not correctly different.

The evidence therefore separates response-space representability from intervention-coordinate assignment. Models can encode intervention geometry when targets are seen, the observed response space has substantial low-dimensional structure, and correct coordinates support held-out reconstruction. IGC emerges when a genuinely unseen intervention must be assigned to a location within that space. This localization motivated a direct test of whether the missing placement information is diffuse or concentrated in a compact, training-derived residual coordinate system.

### Missing unseen-intervention geometry concentrates in a low-dimensional orientation code

To localize the information missing under source-disjoint generalization, we decomposed each held-intervention response into a baseline prediction and coordinates along residual response directions learned from seen interventions only (Fig. 4a). Within each outer source-disjoint fold, the baseline was fit without the held source, and residual axes were learned from out-of-fold residuals among outer-training sources. Only after the basis and all training choices were fixed did we project the true held response onto the first q residual axes. These held-response coefficients are oracle diagnostics of missing information, not deployable predictions. In K562, grouped geometry was -0.110 at q=0, 0.421 at q=1, 0.602 at q=2, 0.762 at q=4 and 0.800 at q=8 (Fig. 4b; Supplementary Fig. 7a). A small number of coordinates therefore recovered a large fraction of the geometry absent from the source-disjoint baseline.

**Figure 4.**
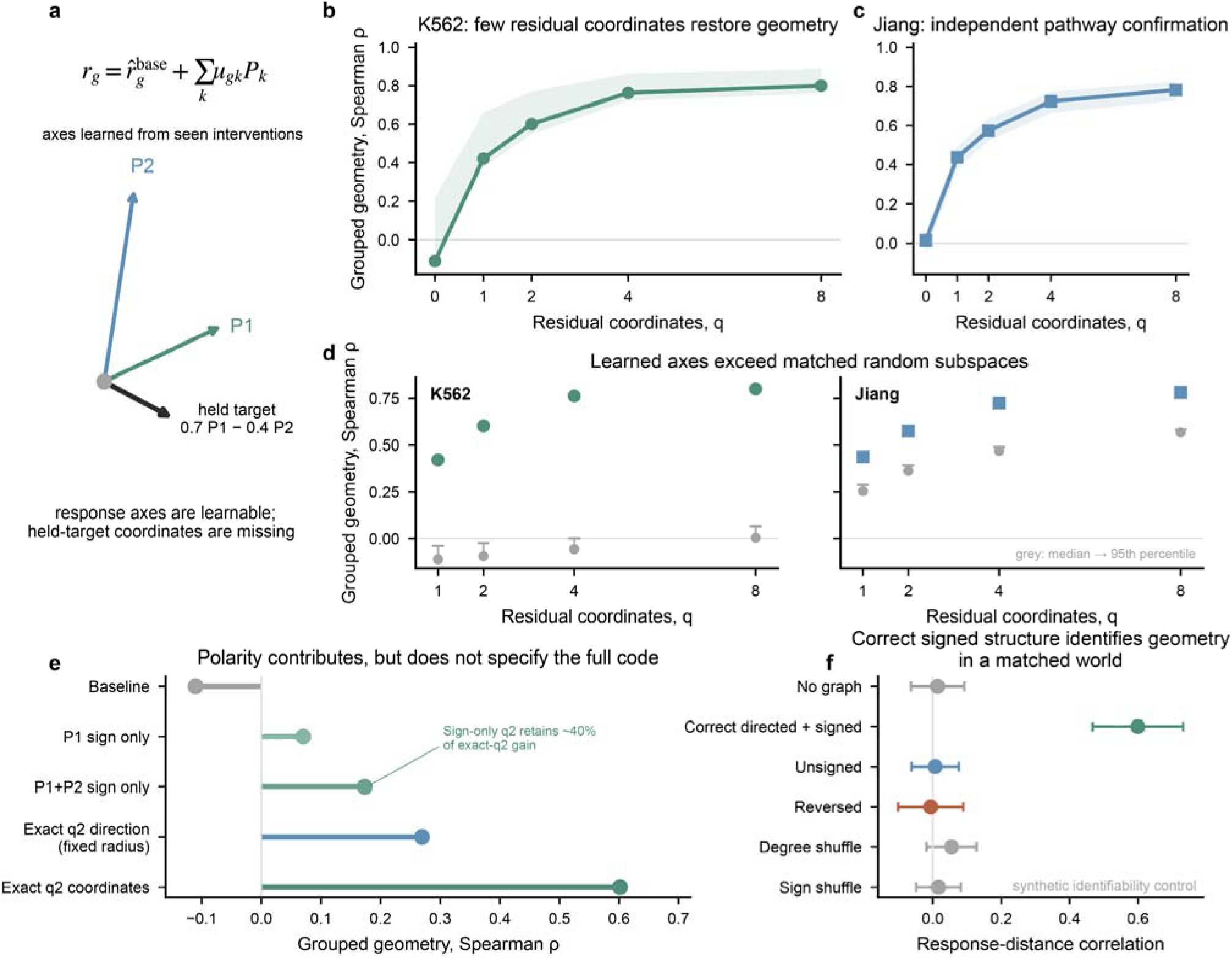
Missing unseen-intervention geometry concentrates in a low-dimensional orientation code. a, Conceptual decomposition of a held response into a source-disjoint baseline and coordinates along residual directions learned from seen interventions; held-target coefficients are oracle diagnostics and are not predicted from held-target covariates. b, K562 grouped geometry as a function of q training-derived residual coordinates; points show aggregate geometry and shading shows the frozen source-bootstrap interval. c, Independent confirmation across IFNB, IFNG and INS perturbations in six Jiang cell lines. d, Learned residual axes compared with 300 matched random orthonormal subspaces for q=1, 2, 4 and 8; grey summaries show the matched null median to 95th percentile and coloured points show the learned axes. Random subspaces receive identical held-response oracle access. e, K562 orientation decomposition comparing baseline, one-sign, two-sign, fixed-radius exact direction and exact continuous q2 coordinates. Sign-only q2 retains 39.86% of the exact-q2 gain, indicating that polarity contributes but does not specify the full code. f, Matched synthetic capacity/identifiability control: correct directed and signed structure identifies unseen geometry in the matched data-generating world, whereas unsigned, reversed, degree-shuffled, sign-shuffled and no-graph controls do not reproduce the rescue. Points and error bars in f show mean ± s.d. across five frozen synthetic worlds. Full orientation-code and sign-decomposition analyses are shown in Supplementary Figs. 7 and 8; real-prior structural controls are shown in Supplementary Fig. 6.

A random-subspace test addressed whether this recovery was simply a generic consequence of low-dimensional held-response projection. For each fold and q, we generated 300 complexity-matched random orthonormal subspaces within the same residual response span and granted them the same diagnostic held-response access as the learned axes. The training-derived axes exceeded the random-subspace 95th percentile for q=1, 2, 4 and 8 in K562 (all empirical P=0.0033; Fig. 4d and Supplementary Fig. 7b). Independent analysis across IFNB, IFNG and INS perturbations in six cell lines from the Jiang dataset^23^ reproduced the pattern: overall grouped geometry increased from 0.013 at q=0 to 0.438, 0.574, 0.724 and 0.782 at q=1, 2, 4 and 8, respectively (Fig. 4c; Supplementary Fig. 7c). Learned axes again exceeded complexity-matched random subspaces at every q (all empirical P=0.0033; Fig. 4d and Supplementary Fig. 7d). These controls identify a reproducible training-derived low-dimensional structure rather than a generic projection advantage.

Polarity captured only part of this coordinate information. Replacing each continuous coordinate by its sign and a training-derived median magnitude improved geometry over baseline but recovered only part of the full signal (Fig. 4e; Supplementary Fig. 8). In K562, the baseline, one-sign, two-sign, fixed-radius exact-direction and exact two-coordinate values were -0.110, 0.070, 0.173, 0.269 and 0.602, respectively. Relative to baseline, the two-sign construction retained 39.86% of the exact two-coordinate gain. Jiang showed the same ordering (0.013, 0.238, 0.316, 0.370 and 0.574). Polarity is therefore informative, but the prespecified strong sign-sufficiency criterion was not met. Magnitudes and continuous coordinate combinations carry additional intervention-specific information.

A synthetic benchmark provided a conditional-identifiability positive control. When the data-generating system and supplied structure were aligned, the correct directed and signed graph identified unseen response geometry. Unsigned, reversed, degree-shuffled, sign-shuffled and no-graph controls did not reproduce the effect (Fig. 4f). This establishes capacity in a world where the structural observables identify intervention orientation; it does not imply that generic biological priors provide the same information in real data. Indeed, OmniPath^21^ and CollecTRI^22^ variants did not robustly rescue real-data geometry (Supplementary Fig. 6). The combined evidence defines the missing variable more precisely: unseen-intervention failure is concentrated in a compact continuous orientation code rather than an inability to represent response space itself.

### Intervention orientation determines trajectory entry and becomes observable after early target perturbation

We used the RENGE iPSC perturbation time course to examine how intervention-specific orientation affects dynamics across Days 2-5 (Fig. 5a). The main analysis focuses on trajectory entry and empirical observability. Detailed first-wave temporal controls, static-breadth comparisons and temporal-order controls are reported in Supplementary Fig. 9; trajectory-entry, oracle-rescue, Markov-formulation and orientation-reliability analyses are reported in Supplementary Fig. 10. These supporting analyses are interpreted locally and do not establish zero-shot identification of a globally unseen source.

**Figure 5.**
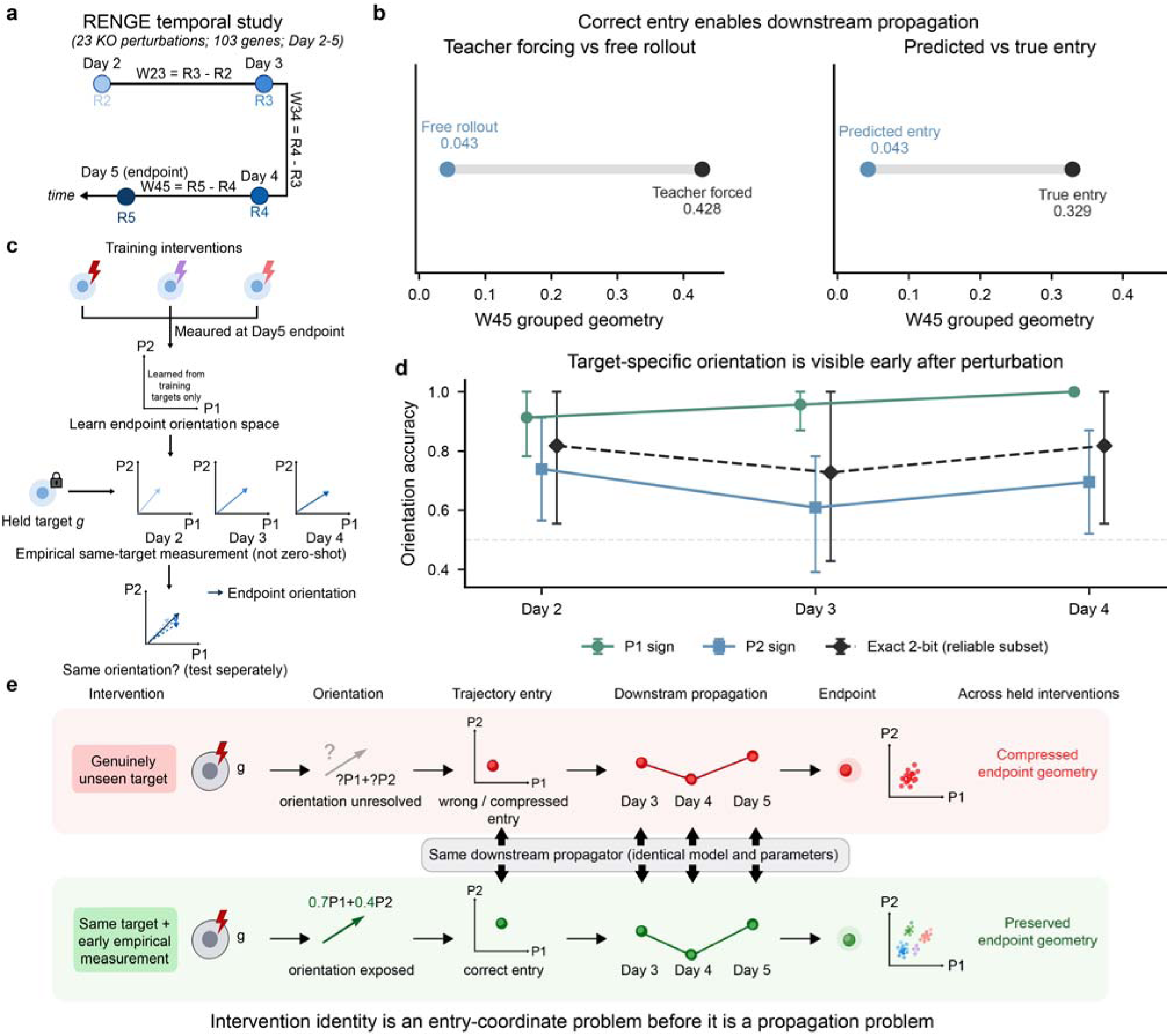
Intervention orientation determines trajectory entry and becomes observable after early target perturbation. a, RENGE temporal design across Days 2-5, with response states R2-R5 and adjacent response waves W23, W34 and W45; R5 is the endpoint state. b, Left, free rollout versus teacher forcing with the true intermediate state (W45 grouped geometry 0.043 versus 0.428); right, predicted versus true entry (0.043 versus 0.329). c, Early target-specific orientation assay. Endpoint P1/P2 axes are learned from training sources only with the held target excluded; the held target is then measured at Day 2, Day 3 or Day 4, with each early time point tested separately and projected onto the same frozen endpoint axes. This is an empirical same-target assay, not zero-shot prediction. d, Accuracy with which Day 2, 3 or 4 held-target responses recover Day 5 orientation. P1 accuracy is 0.913, 0.957 and 1.000; P2 accuracy is 0.739, 0.609 and 0.696. Black diamonds show exact two-axis accuracy among the 11 targets meeting the predefined >=0.8 reliability criterion on both axes; error bars show frozen source-bootstrap intervals. e, Mechanistic synthesis: unresolved orientation yields a misassigned trajectory entry and compressed geometry across held interventions, whereas an early same-target empirical response exposes orientation and permits correct entry into the same downstream propagation process. Supporting temporal controls and orientation-reliability analyses are shown in Supplementary Figs. 9 and 10.

Correcting trajectory entry produced a large rescue of later response geometry. Free rollout yielded W45 grouped geometry of 0.043. Teacher forcing with the true intermediate state raised it to 0.428 (Fig. 5b). In a complementary entry-state comparison, the predicted entry again yielded 0.043, while initialization from the true early entry reached 0.329. Later dynamics therefore become substantially more predictable once the intervention is placed at the correct early state; the dominant error precedes downstream propagation rather than reflecting an inability to propagate a correctly initialized trajectory.

To determine when the missing orientation becomes empirically visible, Day 5 P1/P2 endpoint axes were learned within each outer fold from training sources only, with the held target excluded from baseline fitting and axis construction. The held target’s true Day 2, Day 3 or Day 4 response was then projected separately onto the same fixed axes and compared with its Day 5 orientation (Fig. 5c). P1 sign accuracy was 0.913, 0.957 and 1.000 at Days 2, 3 and 4, respectively. P2 accuracy was lower at 0.739, 0.609 and 0.696 (Fig. 5d). Among the 11 targets meeting the predefined reliability criterion on both axes, exact two-axis sign accuracy was 0.818, 0.727 and 0.818. Split-half analysis confirmed very high P1 reliability (0.990) and more limited P2 reliability (0.684; Supplementary Fig. 10e,f). Because this experiment measures the held target itself, it is an empirical same-target observability assay, not zero-shot prediction.

These temporal results connect the orientation code to trajectory entry. The missing coordinates are not intrinsically unobservable: they become visible rapidly once the target is perturbed. If entry is assigned incorrectly, however, the same downstream propagation machinery can follow a coherent but incorrect trajectory (Fig. 5e). This dynamic localization links the low-dimensional coordinate deficit to the strong benefit of target-specific empirical anchoring tested next.

### Target-specific empirical anchoring resolves intervention identity more efficiently than broad intervention coverage

The orientation-code and trajectory-entry results point to a strongly intervention-specific information deficit. This raises a practical allocation question: for a fixed unseen target, is it more useful to observe many other perturbations or to observe that same target once in another cellular context?

A factorial experiment addressed this question using perturbations shared between K562 and RPE1 cells. A fixed set of 120 target interventions was held out throughout, while coverage of the remaining paired interventions increased from 10% to 90%. In the zero-shot regime, a lightweight Ridge model predicted each target without using an empirical response from that target. In the anchored regime, the measured response of the same target in the source context was provided to a cross-context Ridge map. An identity-shuffled control supplied the response of the wrong target while leaving the fitted map unchanged (Fig. 6a).

**Figure 6.**
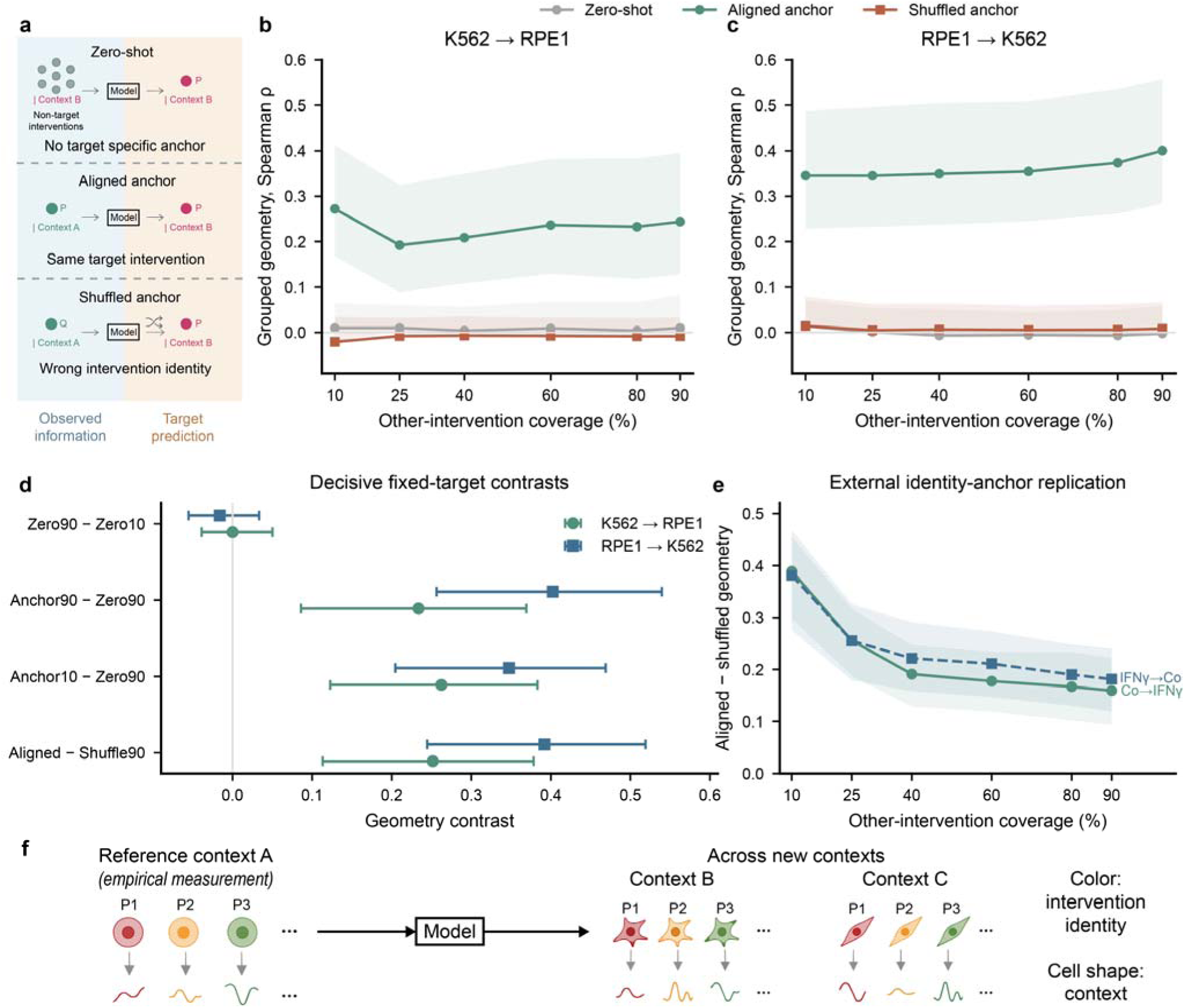
Target-specific empirical anchoring resolves intervention identity more efficiently than broad intervention coverage. a, Information regimes for zero-shot prediction, same-target empirical identity anchoring and an identity-shuffled anchor control. b,c, Nested-coverage factorial experiments for K562→RPE1 and RPE1→K562 using the same 120 fixed held-out targets at every coverage level; lines show five-seed mean grouped geometry and shaded regions show paired target-bootstrap 95% confidence intervals. d, Prespecified fixed-target contrasts comparing increased non-target breadth, aligned anchoring, low-coverage anchoring versus high-coverage zero-shot prediction, and aligned versus shuffled anchors; error bars show paired target-bootstrap 95% confidence intervals. e, External Frangieh Co-culture↔IFNγ replication of the aligned-minus-shuffled anchor advantage across coverage levels, with 95% confidence bands. f, Atlas-design implication in which intervention identity is empirically anchored in a reference context and models generalize those anchored effects across new cellular contexts. The empirical anchor is interpreted as target-specific identity information consistent with the orientation-code mechanism, not as a direct measurement of the exact residual coefficient vector. Internal seed-level and external high-cell-count robustness analyses are shown in Supplementary Fig. 11.

Increasing exposure to other interventions produced essentially no geometric rescue. From 10% to 90% coverage, zero-shot geometry changed by -0.0000 in K562-to-RPE1 transfer (95% confidence interval, -0.0389 to 0.0501) and by -0.0161 in RPE1-to-K562 transfer (-0.0553 to 0.0334; Fig. 6b,c). In contrast, providing the same target’s empirical response in the source context produced large gains. At 90% coverage, aligned anchoring improved geometry over zero-shot prediction by 0.2336 (0.0860-0.3692) for K562-to-RPE1 and by 0.4024 (0.2564-0.5396) for RPE1-to-K562.

The information-efficiency gap was even larger when low-coverage anchoring was compared with high-coverage zero-shot extrapolation. An aligned anchor trained with only 10% paired-intervention coverage still exceeded 90% zero-shot prediction by 0.2624 (0.1227-0.3831) and 0.3474 (0.2045-0.4691) in the two transfer directions, respectively (Fig. 6d). In this setting, observing the same target intervention in one source context was more informative than observing nearly all other available interventions.

The rescue depended on intervention identity. At 90% coverage, aligned anchoring exceeded the identity-shuffled control by 0.2516 (0.1132-0.3783) in K562-to-RPE1 and by 0.3918 (0.2446-0.5192) in RPE1-to-K562. The gain therefore cannot be explained simply by adding another expression vector or generic source-context information. Seed-level coverage trajectories and per-seed decisive contrasts for both transfer directions are shown in Supplementary Fig. 11.

Independent replication used RNA-only processed Frangieh Perturb-CITE-seq data^20^. We analysed 237 interventions shared between Co-culture and IFNγ contexts, with 120 fixed held-out targets and 3,423 strict-trans response genes. Identity-aligned source responses outperformed identity-shuffled responses at every training-coverage level in both directions (Fig. 6e). At 90% coverage, the aligned-minus-shuffled geometry difference was 0.1594 (0.0941-0.2227) for Co-culture-to-IFNγ and 0.1815 (0.1194-0.2400) for the reverse direction. The effect remained positive after restricting analysis to high-cell-count targets (Supplementary Fig. 11). A matched external zero-shot branch was not accessible because the pre-existing safe intervention descriptor covered too few eligible perturbations. We therefore interpret this experiment as a direct external replication of identity-specific anchoring and a partial replication of the full generalization-axis asymmetry.

The experiments reveal a pronounced asymmetry in information efficiency. Broad exposure to non-target interventions provided little improvement for fixed-target zero-shot geometry. A single empirical observation of the same intervention supplied a large and transferable identity anchor. This asymmetry is consistent with the mechanism above: additional interventions can improve knowledge of shared response structure, but they do not directly provide the held target’s orientation coordinates. A same-target empirical response supplies intervention-specific placement information. The result motivates a different allocation of measurement and modelling effort in virtual-cell studies (Fig. 6f): empirically anchor intervention identity in reference contexts and use models primarily to generalize those anchored effects across the substantially larger space of cellular contexts. Internal seed-level and external high-cell-count robustness analyses are shown in Supplementary Fig. 11.

## Discussion

Our results identify a fundamental distinction between reconstructing a plausible perturbed state and preserving the biological identity of the intervention that produced it. Across current perturbation-prediction settings, high conventional similarity can coexist with loss of intervention-specific structure. We refer to this phenotype as Intervention Geometry Compression (IGC): unseen perturbations occupy a predicted response landscape with weaker pairwise organization, degraded local neighborhoods, reduced between-intervention variation and fewer effective response directions than the corresponding biological data.

Several lines of evidence argue that IGC is not primarily a lack of output capacity. The same model can represent intervention geometry for seen targets while losing it for unseen targets, and correct response-basis coordinates support accurate held-out reconstruction. The residual analysis sharpens this localization: in both K562 and the independent Jiang pathway resource, much of the missing geometry is recovered by only a few training-derived residual coordinates. Those axes outperform complexity-matched random subspaces given identical diagnostic access. The central missing quantity is therefore not simply the ability to generate diverse responses, but the assignment of a genuinely unseen intervention to a compact continuous orientation within a learnable response space. Polarity contributes to that assignment, yet sign alone does not specify the full code.

Controlled structural simulations show that unseen-intervention assignment can become identifiable when intervention-side information is genuinely predictive of downstream effects. Correct direction and regulatory sign rescue geometry in the synthetic world; altered structural controls do not. Current OmniPath- and CollecTRI-derived priors, however, fail to reproduce this rescue in real data, indicating that the relevant information should not simply be equated with a generic gene-regulatory network.

Zero-shot intervention prediction is therefore conditional on whether the observables available for an unseen source identify its response orientation. Model expressivity alone cannot supply information absent from those observables. Consistent with this distinction, spontaneous expression covariation in unperturbed RPE1 cells captured shared cellular response structure but did not identify the sender-specific residual geometry of unseen perturbations, indicating that abundant observational variation is not equivalent to target-specific causal excitation.

The temporal analyses provide a dynamic interpretation of the same bottleneck. Teacher forcing and true trajectory entry markedly improve later geometry, showing that downstream propagation capacity is present once the early state is correct. Conversely, the held target’s own early perturbational response reveals its endpoint orientation with high accuracy on the reliable dominant axis. This does not make the early-response assay zero-shot; rather, it establishes an observability boundary. Intervention-specific orientation becomes measurable rapidly after the target is perturbed, linking the low-dimensional coordinate deficit to the trajectory-entry failure seen during free rollout.

The generalization-axis experiment reveals the practical consequence of this bottleneck. If target identity could be inferred efficiently from the response patterns of other interventions, increasing intervention coverage should progressively rescue a fixed unseen target set. Instead, expanding non-target coverage from 10% to 90% produced essentially no improvement. Observing the same intervention in another context generated large identity-specific gains. The aligned-anchor effect reproduced bidirectionally in K562 and RPE1 and independently in the Frangieh multi-context system. The two information sources therefore play different roles: broad intervention coverage helps define shared response structure, while a same-target empirical response provides intervention-specific placement information that can be transferred across context.

These conclusions are deliberately narrower than a universal impossibility claim. The synthetic benchmark demonstrates that zero-shot prediction can work when the available structure is sufficiently informative. The orientation code is also continuous rather than a literal two-bit variable: in the authoritative K562 analysis, two signs retain 39.86% of the exact two-coordinate geometry gain but do not satisfy the prespecified strong sign-sufficiency criterion. The Jiang confirmation is restricted to the high-reliability IFNB, IFNG and INS pathways and should not be generalized to every pathway or context. Likewise, the Frangieh experiment directly replicates identity-specific anchoring but does not provide a matched external zero-shot branch. These boundaries support an identifiability interpretation without claiming that future representations cannot improve unseen-intervention extrapolation.

These findings recast the central design problem for virtual cells: not simply how large a model or perturbation atlas should become, but which biological dimensions should be measured experimentally and which should be delegated to model generalization. For a defined library of atomic interventions, intervention identity is comparatively bounded. Cellular context, by contrast, expands combinatorially across cell type, state, developmental stage, disease, environment, treatment and time. Our results therefore support a practical design principle: measure intervention identity densely enough to anchor its biological orientation, then model how those anchored effects transform across contexts. More generally, bounded biological identities may be better measured, while combinatorial biological contexts remain the natural domain for predictive generalization.

## Methods

### Study design

We evaluated perturbation prediction at the level of control-relative response vectors and intervention relationships rather than relying only on absolute perturbed-state similarity. Unless otherwise specified, a perturbation response was defined as the mean expression state of perturbed cells minus the matched control mean within the same biological context. Main-text geometry analyses used source-disjoint or identity-held evaluation so that test interventions were absent from the relevant recipient-context training data. Gene panels, normalization procedures and quality filters were frozen within each analysis and are reported in the corresponding source manifest.

### Perturbation-response representation

For each perturbation *p* in biological context *c*, we represented the perturbation effect as a control-relative response vector *R_p,c_*, defined as the difference between the pseudobulk mean expression of perturbed cells and the matched control mean from the same context. Means were computed after the dataset-specific preprocessing described below. Analyses used frozen source-level pseudobulk responses rather than individual cells unless explicitly noted. To reduce trivial target-gene effects, geometry analyses used frozen strict-trans gene panels defined in the corresponding audit. For the GEARS RPE1 evaluation, the primary strict-trans panel contained 768 genes. Dataset-specific gene panels and preprocessing choices were held fixed across compared conditions.

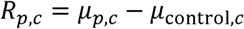

### Conventional state, response and intervention-specific metrics

We evaluated prediction at three progressively more intervention-specific levels. Absolute-state performance compared the predicted perturbed expression state with the observed perturbed state. Response-level performance compared predicted and observed control-relative response vectors. To isolate perturbation identity beyond a response shared across sources, each response was decomposed into the mean response over the fold-reference perturbation set and an intervention-specific residual. The same reference-derived mean response was subtracted from both prediction and observation. The source-ignorant baseline assigned every perturbation this reference mean response and therefore had an identically zero intervention-specific residual.

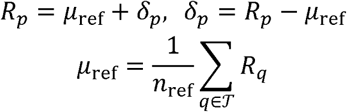

### Intervention geometry

Intervention geometry was defined from pairwise relationships among intervention-specific response vectors. Within each held-out prediction group, response vectors were *L*_2_-normalized and cosine distance was computed for every intervention pair. Global intervention geometry was then defined as the Spearman rank correlation between the predicted and observed vectors of upper-triangular pairwise cosine distances. The metric therefore tests whether the relative ordering of intervention-to-intervention relationships is preserved, independent of a global response-scale change.

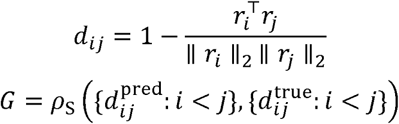

### Artifact-safe grouped out-of-fold evaluation

A central evaluation constraint was that predictions generated by independently trained held-out models were never concatenated into a single response cloud before geometry calculation. Instead, geometry was computed separately within each held-out group using predictions from one trained model, and group-level statistics were then aggregated. This same-model grouped procedure prevents fold-specific offsets or transformations from generating artificial pairwise structure. A dedicated null audit demonstrated the failure mode: naive stitching yielded a geometry value of 1.0 for a source-ignorant null, whereas grouped evaluation returned 0.0. All reported OOF geometry results use the grouped procedure.

### Local geometry, between-source variance and spectral compression

Global geometry was complemented by local and distributional summaries. For each intervention, local neighborhood preservation was quantified as the fraction of its *k* = 10 nearest observed-response neighbors that were also among its 10 nearest predicted-response neighbors, averaged across interventions. A complementary local distance-rank statistic was obtained by computing, for each intervention, the Spearman correlation between its predicted and observed distance vectors to all other interventions in the same held-out group and averaging across interventions. Between-source variance was defined as the mean gene-wise variance across perturbation responses, and variance retention as the predicted-to-observed ratio. Spectral compression was evaluated after centering each response gene across interventions and applying singular-value decomposition. Variance weights were defined from squared singular values and used to compute the PC1 fraction, the number of components required to explain 80%, 90% and 95% of variance, the entropy effective rank and the participation ratio.

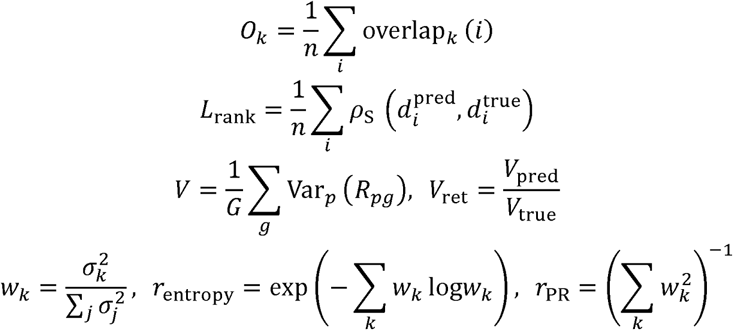

### GEARS and scGPT established-model evaluation

Established perturbation models were evaluated from frozen predictions rather than retrained during manuscript preparation. The GEARS audit used frozen Replogle RPE1 OOF predictions and the same response, strict-trans and grouped-geometry definitions described above. Five independent outer-fold models were used so that held perturbation identities were not used for checkpoint selection in their evaluation fold. The final scGPT audit likewise used five completed frozen OOF folds comprising 1,732 unique held-out sources, with 757 strict-trans genes after excluding eligible perturbation-source genes. Inner training, inner validation and outer OOF sources were disjoint, five distinct best-checkpoint hashes were verified, and outer OOF sources were not used for checkpoint selection. Both established-model audits used the same metric hierarchy, source-ignorant controls, same-model geometry firewall, local geometry, variance-retention and spectral summaries; neither model was retrained for manuscript preparation.

### Seen-versus-unseen capacity analysis

To separate representational capacity from unseen-intervention generalization, we compared intervention geometry for perturbations represented during training with geometry for source-disjoint perturbations using a matched large Transformer analysis in RPE1 cells. The same response construction and geometry implementation were used for both conditions. The comparison was designed to ask whether the architecture can represent intervention structure when source identities are available, rather than to estimate generalization from a mixture of seen and unseen perturbations.

### Response-space PCA and oracle reconstruction

Response-space representability was assessed using a deterministic principal-component analysis of the frozen K562 five-fold training-reference strict-trans response matrix. PCA was fit on the frozen training-side responses for each fold, and cumulative explained variance was summarized across the matched analysis. Across the five training-reference folds, cumulative explained variance reached 0.609, 0.710, 0.797 and 0.878 at 8, 16, 32 and 64 components, respectively. In the oracle reconstruction analysis, the correct held-out response coordinates in the learned basis were supplied and mapped back to gene space; reconstruction performance was evaluated at the same component counts. This oracle analysis tests representability of held-out responses conditional on correct coordinates and is not a deployable prediction experiment. This response-space oracle asks whether held responses can be represented when their global response-basis coordinates are known. It is distinct from the residual orientation-code analysis below, which asks how many coordinates are required to correct a source-disjoint baseline prediction.

### Source-side identifiability analysis

We directly tested whether an intervention representation constructed from unperturbed control cells could identify the geometry of subsequent perturbation-specific responses. For each measured source gene, EstablishedOBS71 concatenated seven control-cell expression summaries (mean, standard deviation, detection frequency, 25th, 50th and 75th percentiles, and the log-transformed number of expressing cells) with 64 components obtained by truncated singular-value decomposition of standardized gene-by-control-cell expression profiles. Within each of five source-disjoint folds, feature scaling was fitted on reference source genes only and then applied to held-out query genes. Cosine similarity was computed between all pairs of held-out EstablishedOBS71 vectors. Response similarity for the same pairs was computed from fold-reference-mean-centered strict-trans response vectors. Source-side identifiability was quantified as the Spearman correlation between the two pairwise similarity vectors within each fold and summarized across folds. The canonical analysis included 1,755 held-out source evaluations and 307,282 within-fold source pairs.

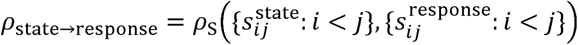

### Natural-fluctuation source-anchor analysis

We tested whether spontaneous expression variation among unperturbed cells provided a target-specific response-space direction for source-disjoint perturbations. The analysis used the processed Replogle RPE1 dataset containing 240,774 cells and 8,749 measured genes; 11,485 cells annotated as perturbation == "control" were treated as the unperturbed population. The canonical five source-disjoint folds were reused without modification (350, 361, 352, 337 and 355 held-out targets), yielding 1,755 held-out target evaluations. Primary response-space analyses used the 768 strict-trans genes shared across all five folds, with perturbation-source genes excluded from the response panel.

For each eligible target gene, we constructed control-derived response-space vectors across strict-trans genes using the target’s sample covariance, Pearson correlation and univariate response-on-target slope with each response gene across unperturbed cells. Because the perturbations were inhibitory, each association vector was multiplied by −1 to orient it according to the nominal decrease of the target. The processed H5AD X matrix was used as the existing nonnegative normalized/log-transformed expression matrix; no raw or count layer was substituted. For residualized estimators, the full 11,485 × 8,749 control-cell matrix was densified for scikit-learn PCA with 20 components, the randomized solver and random state 1701. A library proxy was defined as each cell’s row sum in the existing normalized/log-transformed matrix. The library proxy and PC1–PC20 were standardized, and a single rank-22 design matrix [intercept, standardized library proxy, standardized PC1–PC20] was fit jointly by ordinary least squares using the Moore–Penrose pseudoinverse to the target and response columns. Negative target–response covariance and correlation were then recomputed from the residuals, with covariance divided by the number of control cells minus the design rank. All control-derived vectors used control cells only and never cells carrying the corresponding held-out perturbation.

Natural-fluctuation vectors were evaluated against both the full control-relative response and the fold-reference-mean-centered intervention-specific residual. Target-level correspondence was quantified by cosine, Pearson and Spearman similarity together with signed-direction metrics. Target-permutation nulls reassigned fluctuation vectors across source genes using 200 frozen permutations. Held-out truth was used only in explicitly labelled oracle projections, in which the scalar projection of the observed response onto a control-derived direction defined an information ceiling. Deployable zero-shot calibration instead fitted a single global scalar using reference-fold targets only and applied it unchanged to held-out targets.

Intervention geometry was evaluated using the same source-disjoint, strict-trans and within-fold grouped-geometry firewall used throughout the study. We quantified grouped pairwise geometry, k-nearest-neighbour overlap, target retrieval, between-intervention variance retention and spectral summaries. Primary inference emphasized the intervention-specific residual because correspondence with the full control-relative response can be driven by programs shared across perturbations. The analysis used base random seed 20260818, retained the frozen fold seed 1701 and used PCA seed 1701.

#### Low-dimensional residual orientation-code analysis

All analyses used outer source-disjoint folds. Within each outer fold, held intervention sources were excluded from baseline fitting, residual-axis construction and hyperparameter selection. A frozen receiver/incoming-based Ridge baseline was fitted using dataset-specific source-side features; regularization was selected only among outer-training sources. To prevent in-sample residual leakage, predictions for outer-training sources were generated by inner source-disjoint models that excluded the corresponding source. The outer-training residual matrix was defined as observed response minus these out-of-fold predictions, and the residual basis was obtained by singular-value decomposition of that training residual matrix. Conceptually, the held response is decomposed as follows:

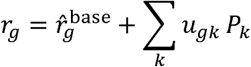

After the residual basis was frozen, the oracle coefficient of held source g on axis k was defined only for diagnostic evaluation as:

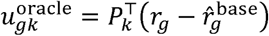

For a held response and its frozen source-disjoint baseline prediction, a q-coordinate oracle reconstruction was obtained by projecting the held residual onto the first q training-derived residual axes:

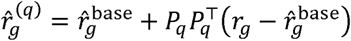

The projection uses the true held response and is therefore an oracle diagnostic used only to localize missing information; it was never used to learn the basis, choose *q or select hyperparameters. We* evaluated *q* = 0,1,2,4,8 using the same grouped, same-model Spearman geometry as elsewhere. K562 used five outer folds over 86 eligible perturbation sources. Displayed aggregate intervals used 2,000 frozen source-bootstrap replicates.

#### Matched random-subspace control

For each outer group and *q* = 0,1,2,4,8, we sampled 300 random orthonormal *q*-dimensional bases within the same training-derived residual span. Each random basis received the identical oracle privilege as the learned basis: the held residual was projected only after the basis and baseline were fixed. Learned-axis geometry was compared with the null median and 95th percentile. The finite-replicate empirical *P* value was calculated as (1 + number of null values at least as large as the observed value)/(1 + number of replicates). Randomization used master seed 20260817 with deterministic child seeds recorded in the frozen manifests.

#### Jiang independent orientation-code confirmation

We analysed the published Jiang pathway Perturb-seq resource23 and its associated differential-expression archive (DE_results_all_pathway.zip). The primary analysis was restricted a priori to IFNB, IFNG and INS, the three pathways that passed the response-geometry reliability audit, across six cell lines (A549, BxPC-3, HAP1, HT-29, K562 and MCF-7). Within each pathway-by-recipient-cell block, held regulators were treated as globally unseen perturbation sources. The frozen baseline used allowed incoming beta-coefficient features from the five auxiliary cell-line contexts, with all baseline fitting, inner validation and residual-axis construction restricted to outer-training sources. Response geometry was scored on the frozen strict-trans log2FC response panel. Before the orientation analysis, pathway reliability was evaluated by deterministic response-gene split halves; IFNB, IFNG and INS had mean geometry reliabilities of 0.896, 0.901 and 0.964, respectively. Aggregate uncertainty used 2,000 bootstrap resamples of pathway-by-cell-line blocks. TGFB1 and TNFA were not used for the primary orientation-code claim because their historical strict-trans geometries were materially less reliable.

#### Sign-only and fixed-radius orientation decomposition

For each residual axis k, the training-derived reference magnitude was defined as:

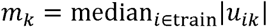

The sign-only held-source reconstruction was:

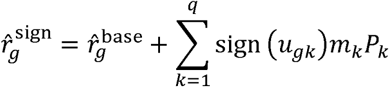

We compared the source-disjoint baseline, one-axis sign reconstruction, two-axis sign reconstruction, exact two-dimensional direction rescaled to a training-only fixed radius, and exact continuous two-coordinate oracle reconstruction. For the two-axis comparison, the retained fraction of the exact-coordinate geometry gain was defined as:

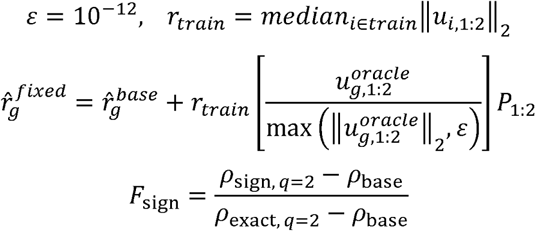

Under the frozen K562 construction, *F_sign_* was 0.3986 (39.86%). The prespecified strong sign-sufficiency gate was not passed; sign-based results are therefore interpreted as evidence that polarity contributes information, not that it fully specifies unseen intervention identity.

#### Early target-specific orientation observability

The RENGE orientation assay used the same 23 perturbation sources measured on Days 2-5. Sources were assigned to four frozen outer folds. For each fold, the endpoint baseline and residual *P*_1_/*P*_2_ axes were learned from outer-training sources only, and all 23 perturbation-source genes were excluded from the fixed 80-gene trans-response panel before axis construction. The held target was excluded while the Day 5 endpoint basis was learned. After freezing the basis, the true Day 2, Day 3 or Day 4 response of that same target was baseline-adjusted and projected separately onto *P*_1_/*P*_2_; no combination of multiple early time points was required. The signs of these early coefficients were compared with the held target’s Day 5 coefficient signs. We report per-axis accuracy, balanced accuracy and exact two-axis sign accuracy with 2,000 source-bootstrap intervals. Because the held target is experimentally observed at the early time point, this procedure is an empirical same-target observability diagnostic rather than a zero-shot prediction task.

#### Orientation-sign reliability

Endpoint sign reliability was evaluated before interpreting the early-orientation assay. For each target and outer fold, Day 5 cells were randomly divided into two halves 500 times using deterministic target/fold seeds. Each half-pseudobulk response was baseline-adjusted and projected onto the frozen outer-training *P*_1_/*P*_2_ axes. Aggregate half-versus-half sign reliability was 0.990 for *P*_1_ and 0.684 for *P*_2_. Reliability-qualified exact-state analyses required per-target reliability of at least 0.8 on both axes, yielding 11 targets. This predefined subset was used only to qualify the exact two-axis analysis and did not alter the all-target *P*_1_ or *P*_2_ results.

### Synthetic directed and signed structural benchmark

To test whether intervention geometry is learnable when intervention-to-effect structure is known, we used a controlled synthetic generator with directed and signed relationships. The positive-control condition supplied structure aligned with the generator used to produce the responses. Matched controls removed or altered components of that structure, including unsigned, reversed, degree-shuffled, sign-shuffled and no-graph conditions. This experiment is interpreted strictly as a capacity and conditional-identifiability positive control: it establishes that correctly aligned predictive structure can identify unseen geometry in a matched world where the relevant intervention-to-effect mapping is informative by construction. It is not evidence that the same structure is available from current generic real biological priors.

### Real biological prior controls

We separately evaluated real directed biological priors, including OmniPath- and CollecTRI-derived routing representations. Outgoing, incoming, signed, unsigned, reversed and shuffled variants were compared under the frozen unseen-intervention evaluation. The purpose was to test whether generic real-world network priors provide target-specific predictive direction sufficient to reproduce the synthetic rescue. These analyses define a real-prior boundary and are reported in Supplementary Fig. 6; because the prespecified directional gate was not met, they are not treated as a main-paper solution to IGC.

### RENGE time-resolved perturbation data

Temporal analyses used the RENGE human iPSC CRISPR dataset (GEO GSE213069), comprising 14,945 cells, 103 measured genes and 23 knockout perturbations observed at Days 2, 3, 4 and 5. For perturbation *p* at day *t*, the response was defined as the perturbation pseudobulk mean minus the matched control mean at the same time point:

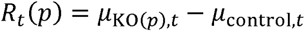

Consecutive response waves were defined by:

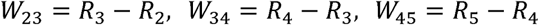

The Day-5 response *R*_5_ was treated as the endpoint state and was not equated with *W*_45_.

### Temporal propagation, teacher forcing and trajectory-entry tests

We first characterized first-wave temporal predictability using the frozen CorrectLag analysis and matched static, same-wave, temporal-shuffle, gene-identity-shuffle and intermediate-masked controls (Supplementary Fig. 9). To separate propagation from trajectory entry, the primary temporal contrasts compared teacher-forced prediction, in which a true intermediate wave was supplied to the frozen propagator, with free rollout initialized from a predicted unseen early response, and separately initialized later propagation from either the true *W*_23_ response or the predicted *W*_23_ response. Oracle direction and program-component substitutions were used as supporting autopsies to determine whether entry errors reflected response magnitude, direction or program composition (Supplementary Fig. 10).

### Endpoint information and Markov autopsy

Endpoint analyses quantified how much Day 5 intervention geometry became recoverable as true temporal information was supplied. Supporting Markov-autopsy experiments tested direct endpoint prediction, impulse propagation, persistent additive and conditional source forcing, additional temporal history and stage-specific dynamics. These variants were evaluated for deployable rollout and for the relationship between response geometry and between-intervention variance (Supplementary Fig. 10). They are used to localize the trajectory-entry bottleneck and are not interpreted as evidence that temporal information from other interventions identifies the orientation of a globally unseen source.

### Internal generalization-axis factorial experiment

We tested the relative information value of broad non-target intervention coverage and target-specific empirical anchoring using perturbations paired between K562 and RPE1. A fixed set of 120 target perturbations was sealed for evaluation. The remaining paired perturbations were subsampled into nested training sets corresponding to 10%, 25%, 40%, 60%, 80% and 90% coverage, using five frozen subsampling seeds. The same fixed targets were used for all coverage levels, seeds and transfer directions. All mappings used lightweight multi-output Ridge regression with a fixed regularization setting specified in the frozen configuration and without target-dependent hyperparameter tuning.

### Frangieh external replication

External replication used the processed RNA-only Perturb-CITE-seq melanoma-TIL dataset, with Control, IFNγ and Co-culture contexts identified from the supplied metadata. The primary transfer pair, selected before performance evaluation on the basis of shared intervention coverage and data completeness, was Co-culture and IFNγ. The analysis contained 218,331 cells, 237 eligible shared gene-level interventions, 120 fixed held targets, 117 non-target training interventions and 3,423 frozen strict-trans response genes. Within each context, counts were normalized by CP10K followed by log1p transformation, pseudobulk means were computed for each perturbation, and the matched context-control mean was subtracted to form response vectors.

### External anchor and identity-shuffle evaluation

For each direction of the Frangieh transfer, multi-output Ridge maps were fit from source-context responses to recipient-context responses using nested 10%, 25%, 40%, 60%, 80% and 90% subsets of the 117 non-target paired interventions. The aligned condition supplied the source-context response of the same held target; the shuffled condition reused the identical fitted map but supplied a deterministically mismatched target response. The external zero-shot branch was not run because the pre-existing safe intervention descriptor covered only 39 of 237 eligible interventions, below the frozen minimum of 60 required for that branch. No new gene representation was created after observing the internal result. A predefined high-cell-count target analysis was used as a robustness check against differences in pseudobulk precision.

### Uncertainty estimation and statistical comparisons

Primary uncertainty for geometry contrasts was estimated by paired bootstrap resampling of held perturbation identities so that compared regimes were evaluated on the same resampled targets. The internal generalization factorial and Frangieh replication used 500 paired bootstrap iterations. Reported intervals are 95% bootstrap confidence intervals. Coverage sets, held targets, shuffle definitions and model settings were frozen before evaluation. For multi-seed coverage analyses, the manuscript reports the frozen seed summaries together with paired target-level uncertainty. No multiplicity correction was used to convert exploratory comparisons into significance claims.

### Software, provenance and reproducibility

Core deterministic audits were performed in Python with NumPy, SciPy and scikit-learn. The repository contains locked GEARS and scGPT environment specifications and an environment manifest for external-model provenance. GEARS and scGPT predictions/checkpoints are treated as frozen expensive assets and are not retrained to regenerate manuscript tables or figures. The geometry firewall, GEARS audit, PCA response-basis analysis, residual orientation-code revalidation, RENGE temporal analyses, generalization factorial, Frangieh replication and natural-fluctuation source-anchor analysis each have frozen scripts/configurations and repository-relative provenance records. The natural-fluctuation release records the processed RPE1 H5AD checksum and exports derived source tables that regenerate Supplementary Fig. 5e–h without loading or redistributing the third-party H5AD. Deterministic CPU replays range from minutes to tens of minutes; the internal factorial required approximately 3 min and the Frangieh replication approximately 2 min once the 1.46-GB processed dataset was local. Public dataset provenance and hashes are recorded in the corresponding data manifests.

## Data availability

The study analyses publicly available single-cell perturbation datasets, including Replogle Perturb-seq, the RENGE time-resolved CRISPR dataset (GEO accession GSE213069), Frangieh Perturb-CITE-seq, and the Jiang pathway Perturb-seq resource. The Jiang differential-expression archive used here is available from the authors’ associated Zenodo dataset (DOI: 10.5281/zenodo.14518762). Data generated from this study is available on Zenodo DOI: <u>10.5281/zenodo.22005137</u>.

## Code availability

The custom code used to reproduce the analyses and figures reported in this study is publicly available at GitHub at https://github.com/yongqih/virtual-cell-intervention-geometry. The final audited release, including the frozen analysis configurations, reproducibility manifests, figure-generation scripts and validation utilities used in this study are available on Zenodo DOI: <u>10.5281/zenodo.22005122</u>. The code is distributed under the MIT License.

## Acknowledgements

We thank members of the Wilson laboratory for helpful discussions and feedback on the study. We also thank the developers and maintainers of the public perturbation datasets and software resources used in this work.

## Author contributions

Y.H. conceived the study, designed and implemented the computational analyses, performed the statistical analyses, generated the figures and wrote the original manuscript. H.W. contributed to methodological discussion and interpretation of results. P.W. supervised the study, provided scientific guidance, contributed to interpretation of the results and revised the manuscript. All authors reviewed and approved the final manuscript.

## AI-assisted tools

Generative AI tools (OpenAI ChatGPT and Codex) were used to assist with code drafting and debugging, methodological brainstorming and language editing. All analytical decisions, statistical evaluations, interpretation of results and final manuscript content were reviewed and verified by the authors.

