## Supplemental Material for "Virtual-cell models compress unseen intervention geometry through a target-specific generalization bottleneck"

This Supplementary Information contains Supplementary Figures 1–11. Exact figure source data and provenance are provided in the accompanying data-availability package.

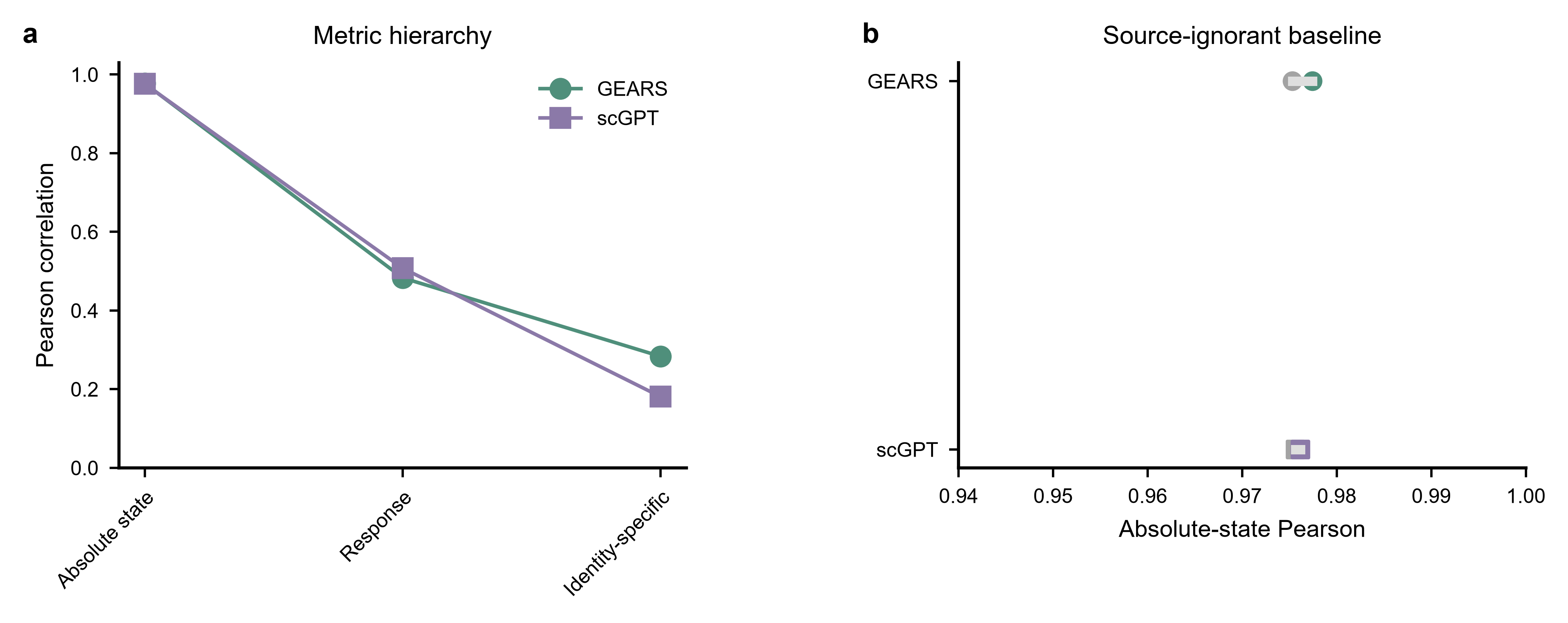

**Supplementary Figure 1 | Metric decomposition and source-ignorant controls.** a, GEARS and scGPT metric hierarchy across absolute perturbed state, total control-relative response and intervention-specific residual. b, Absolute-state Pearson correlation for each learned model and its matched source-ignorant mean-response baseline. The source-ignorant predictor uses no perturbation identity and has an identically zero intervention-specific residual, demonstrating why strong state-level similarity is not sufficient evidence of intervention-specific prediction.

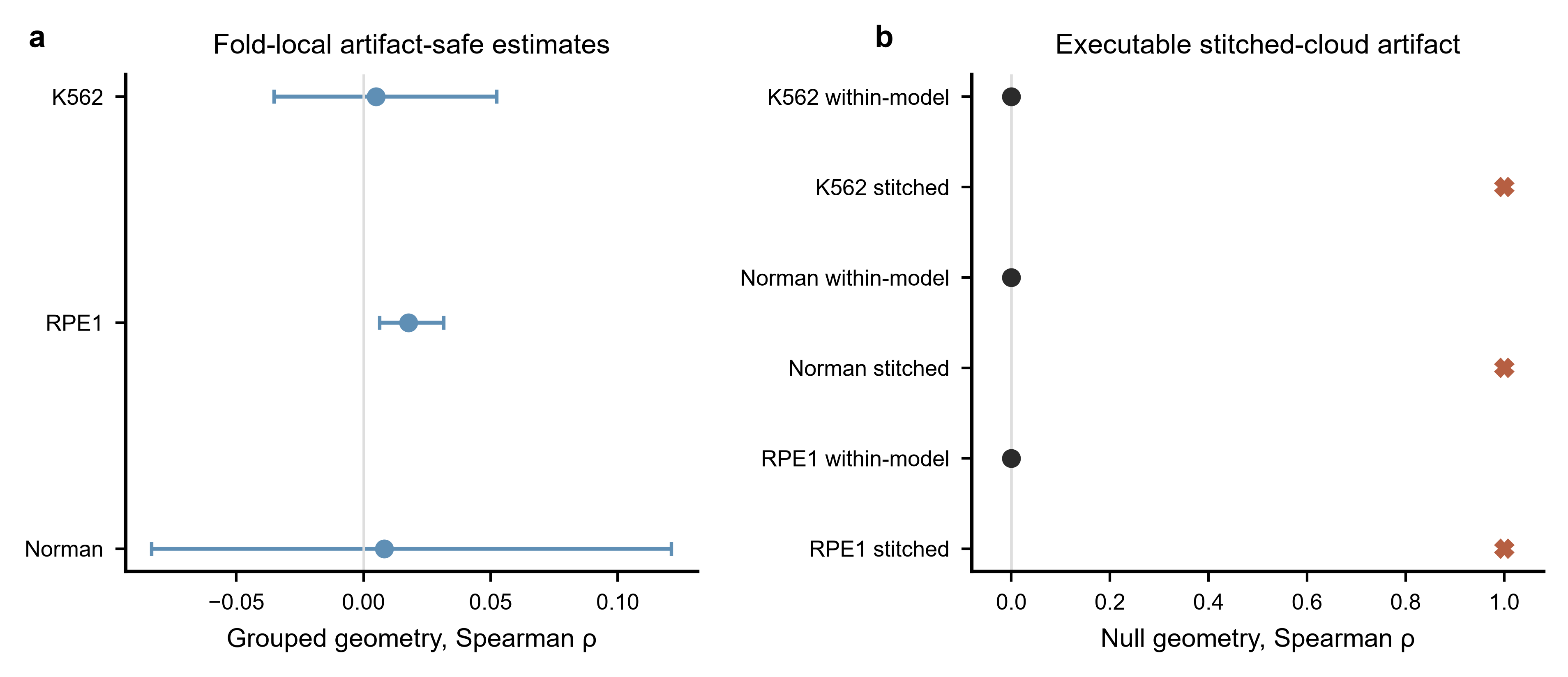

**Supplementary Figure 2 | Artifact-safe intervention geometry.** a, Fold-local same-model OOF geometry estimates for K562, RPE1 and Norman perturbation settings. b, Executable null demonstrating the cross-model stitching artifact: within-model geometry remains near zero for a source-ignorant null, whereas concatenating predictions from independently fitted models can manufacture an apparent geometry approaching 1. All main-text OOF geometry therefore uses same-model grouped evaluation.

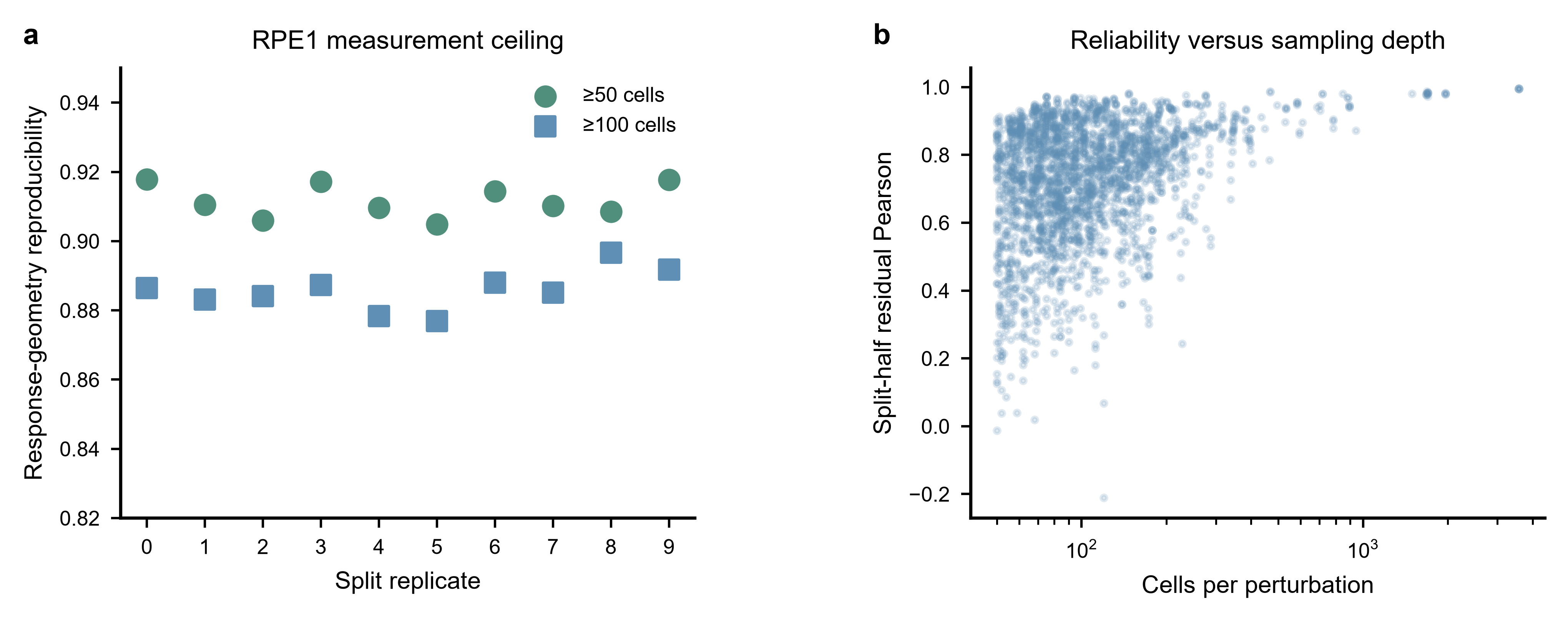

**Supplementary Figure 3 | Response-geometry measurement reliability.** a, Split-cell pseudoreplicate response-geometry reproducibility in RPE1 for perturbations with at least 50 or 100 cells. b, Split-half residual-response Pearson correlation as a function of cells per perturbation. These analyses quantify the measurement ceiling of the observed response geometry and show that the compression phenotype is not explained solely by unstable biological targets.

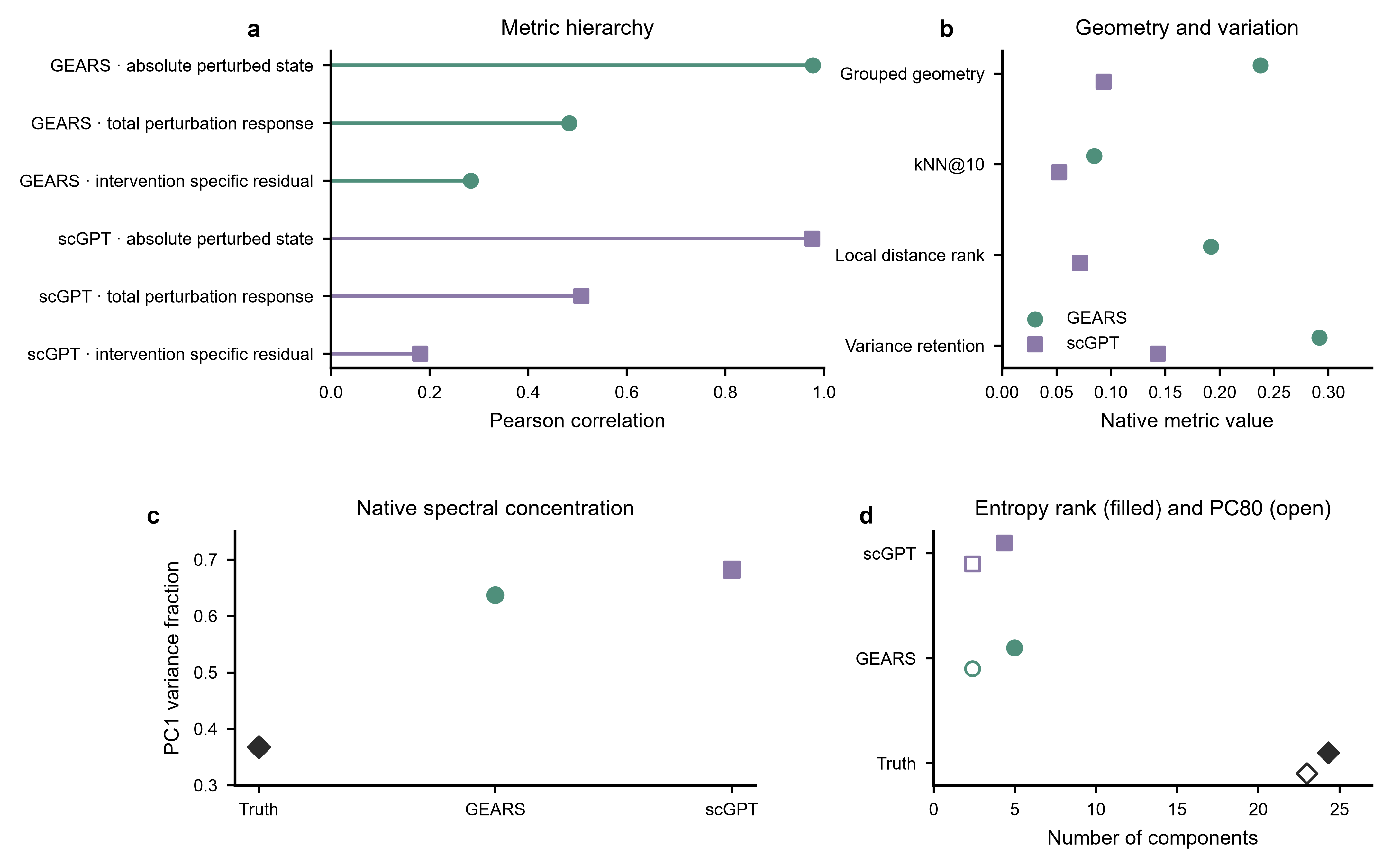

**Supplementary Figure 4 | Full established-model geometry audit.** a, Absolute-state, total-response and intervention-specific residual Pearson correlations for GEARS and scGPT. b, Native grouped geometry, kNN@10 overlap, local distance-rank preservation and between-intervention variance retention. c, PC1 variance fraction for matched truth and model predictions. d, Entropy effective rank (filled markers) and PC80 (open markers) for Truth, GEARS and scGPT. The same source-disjoint, strict-trans and same-model geometry definitions are used throughout.

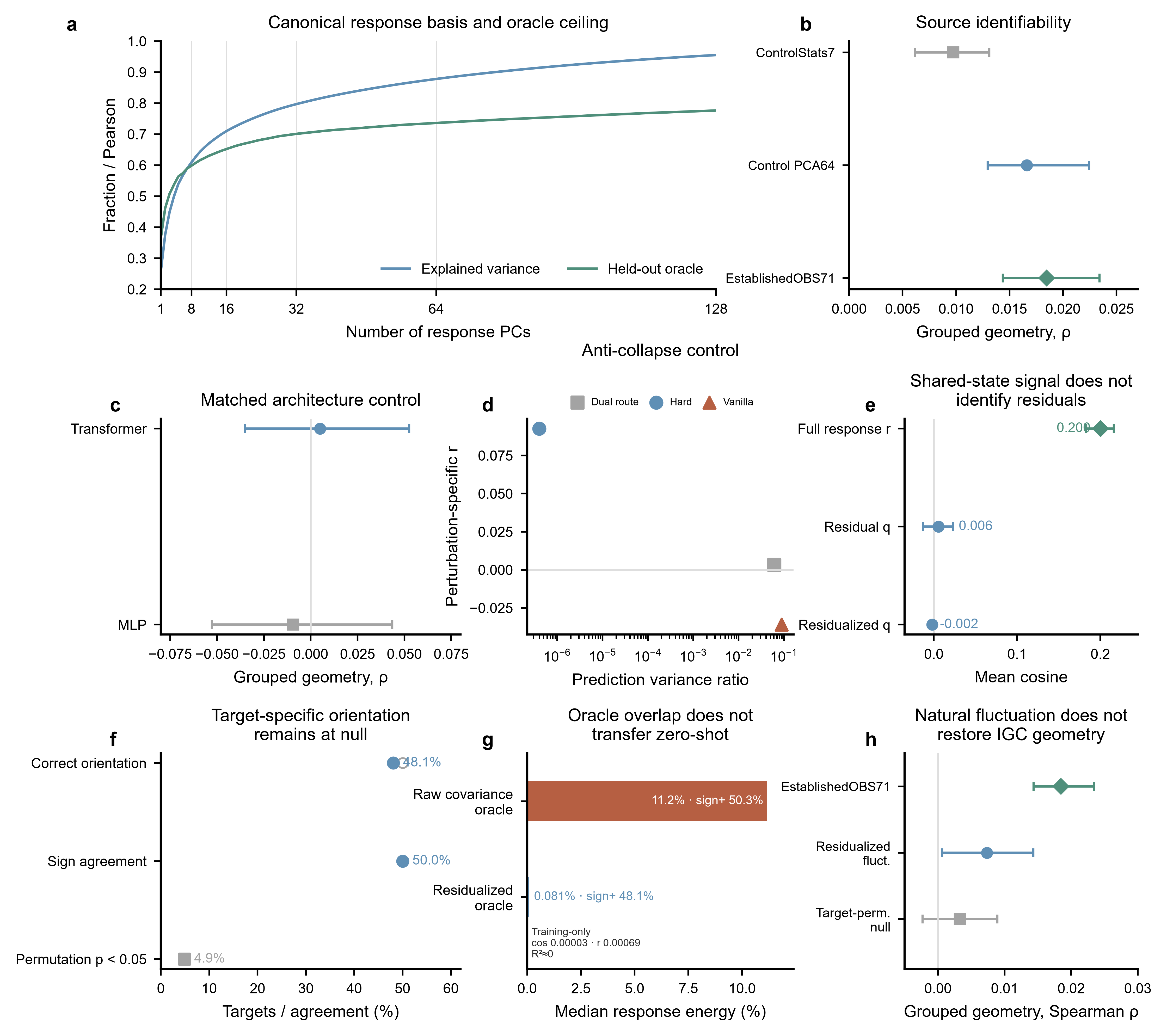

**Supplementary Figure 5 | Capacity, response basis and source identifiability.** a, Canonical response-space PCA cumulative variance and held-out oracle reconstruction as the number of response components increases; guides highlight the prespecified 8, 16, 32 and 64 component levels. b, Source-side identifiability for three leakage-safe control-state representations, including EstablishedOBS71. c, Matched Transformer and MLP grouped geometry. d, Anti-collapse variants showing that changes in prediction variance do not reliably restore perturbation-specific geometry. e, Natural control-cell fluctuation vectors showed apparent correspondence with the full control-relative response, but this signal disappeared for the intervention-specific residual and after residualization of major cell-state variation. f, Target-specific fluctuation orientation remained at null, with directional and sign agreement near chance and only the null-expected fraction of targets exceeding the target-permutation benchmark. g, An unconstrained held-response oracle captured limited response energy from raw covariance only by using the true response to determine target-specific projection sign; this apparent overlap disappeared after state residualization and did not transfer to training-only zero-shot prediction. h, Fluctuation-derived predictions failed to restore held-out intervention geometry and remained below the existing EstablishedOBS71 source descriptor, with negligible between-intervention variance retention. Together, these controls separate response-space representability and shared observational variation from the target-specific information required to assign unseen interventions to the correct response geometry.

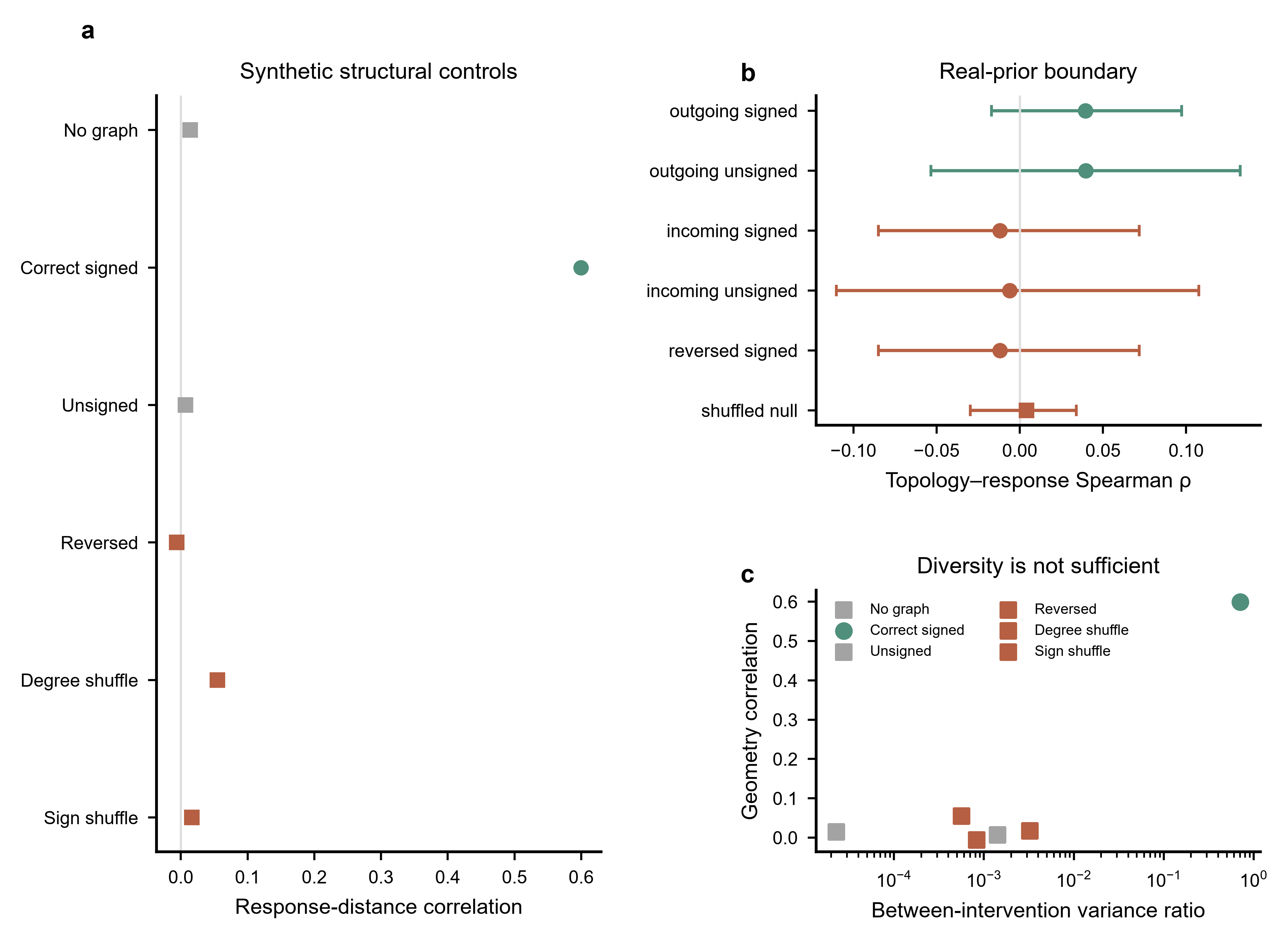

**Supplementary Figure 6 | Structural rescue and real-prior boundary.** a, Complete synthetic structural-control series comparing no graph, correctly directed-and-signed structure, unsigned, reversed, degree-shuffled and sign-shuffled conditions. b, Real OmniPath/CollecTRI-derived outgoing, incoming, signed, unsigned, reversed and shuffled-prior controls. c, Between-intervention variance ratio versus geometry correlation across synthetic structural conditions. Correct generator-aligned structure rescues geometry in the controlled system, whereas the tested generic real priors do not provide a robust real-data rescue.

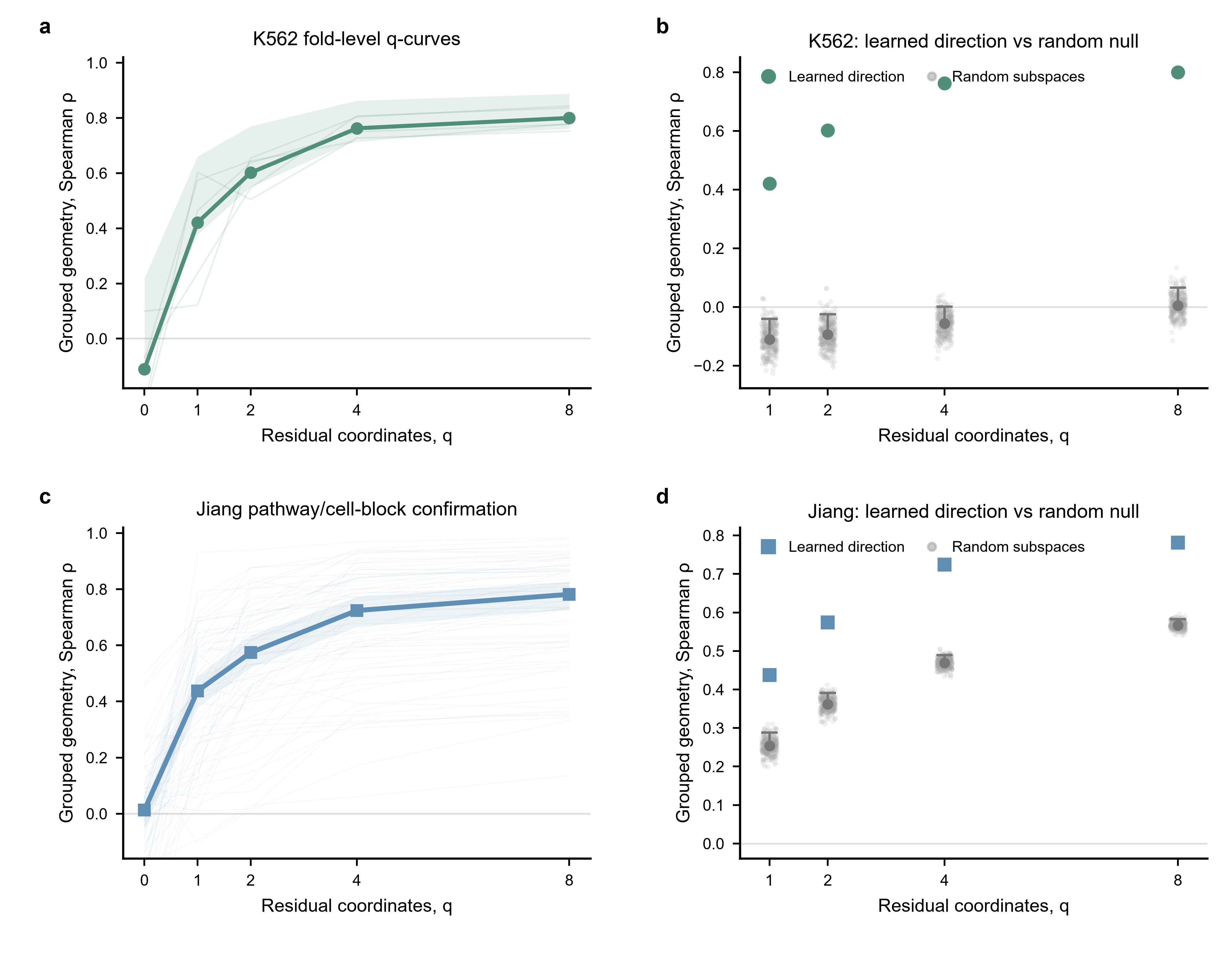

**Supplementary Figure 7 | Low-dimensional orientation-code revalidation and specificity.** a, K562 outer-fold q-curves (thin lines) and frozen aggregate (bold line and interval). b, K562 training-derived residual-axis geometry compared with 300 matched random q-dimensional subspaces at q=1, 2, 4 and 8; grey points show null replicates and grey summaries show the null distribution, while coloured points show the learned directions. c, Jiang IFNB/IFNG/INS pathway-by-cell-line block curves (thin lines) and aggregate confirmation. d, Jiang learned directions versus matched random-subspace nulls. All residual bases are learned from outer-training sources only, and random subspaces receive identical oracle access to held responses.

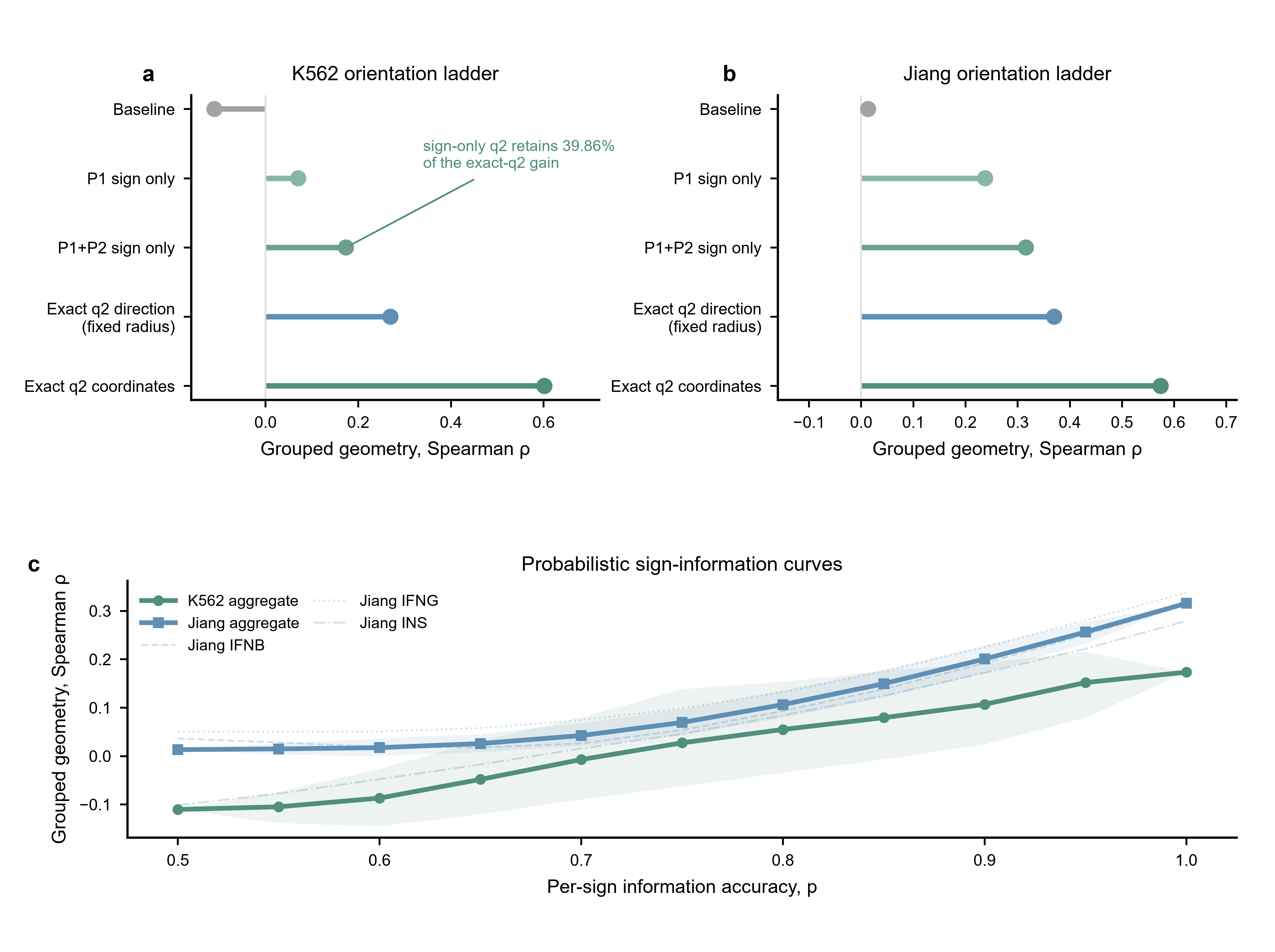

**Supplementary Figure 8 | Orientation decomposition shows that sign is informative but incomplete.** a,b, K562 and Jiang orientation ladders comparing the source-disjoint baseline, one-sign reconstruction, two-sign reconstruction, exact two-dimensional direction at a training-only fixed radius and exact continuous q2 coordinates. In K562, sign-only q2 retains 39.86% of the exact-q2 gain. c, Frozen probabilistic sign-information curves. The K562 and Jiang aggregate curves show how geometry changes as simulated per-sign information accuracy increases; lighter Jiang curves show IFNB, IFNG and INS separately. These analyses do not establish sign sufficiency or a universal accuracy threshold; continuous coordinates remain more informative.

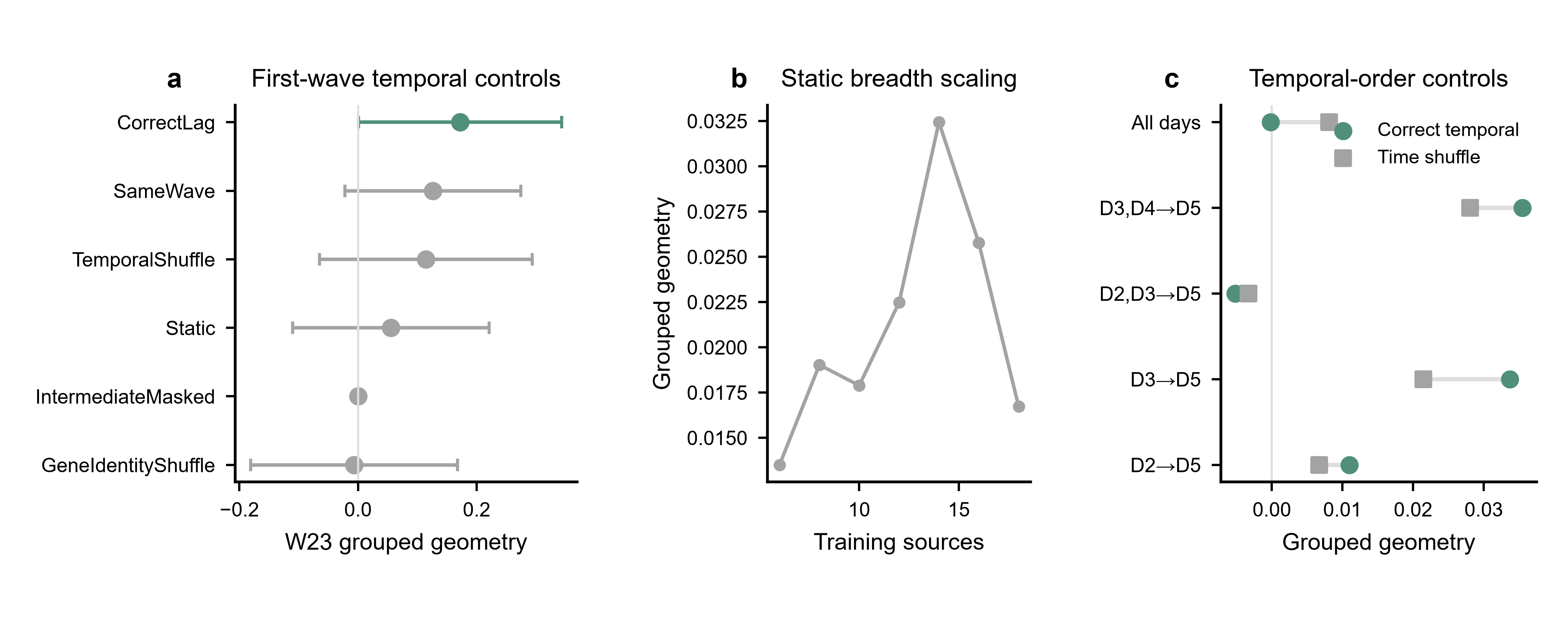

**Supplementary Figure 9 | Temporal signal and first-wave controls.** a, RENGE first-wave grouped geometry for CorrectLag, SameWave, TemporalShuffle, Static, IntermediateMasked and GeneIdentityShuffle controls. b, Static training-source breadth scaling. c, Temporal-order controls comparing correct temporal order with time-shuffled data across alternative day combinations. These analyses characterize local first-wave predictability and the dependence of temporal effects on lag definition and observation window; they do not establish source-disjoint identification of the orientation of a globally unseen intervention.

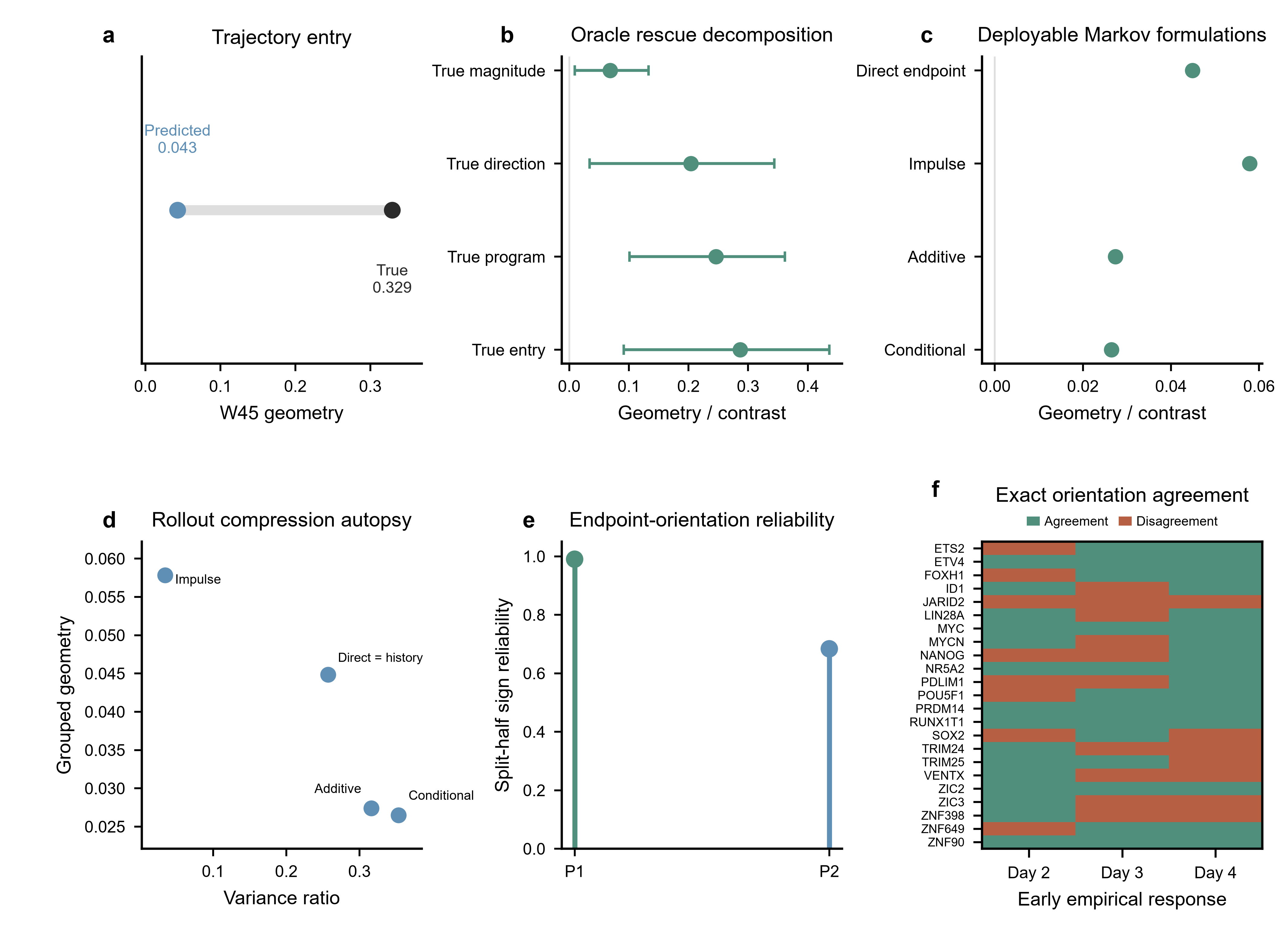

**Supplementary Figure 10 | Trajectory entry, formulation autopsy and orientation reliability.** a, Later-wave geometry after predicted versus true W23 trajectory entry. b, Oracle rescue decomposition replacing response magnitude, direction, program composition or the complete entry state with the corresponding true quantity; error bars show frozen confidence intervals. c, Fully deployable R5 geometry for direct endpoint, impulse, additive and conditional formulations. d, Rollout-compression autopsy relating endpoint geometry to between-intervention variance under alternative formulations. e, Day 5 split-half endpoint-orientation sign reliability on frozen training-derived axes (P1=0.990; P2=0.684). f, Per-target exact early-to-endpoint orientation agreement for Days 2–4; green indicates agreement and rust indicates disagreement. Reliability analyses precede and qualify the early-orientation interpretation.

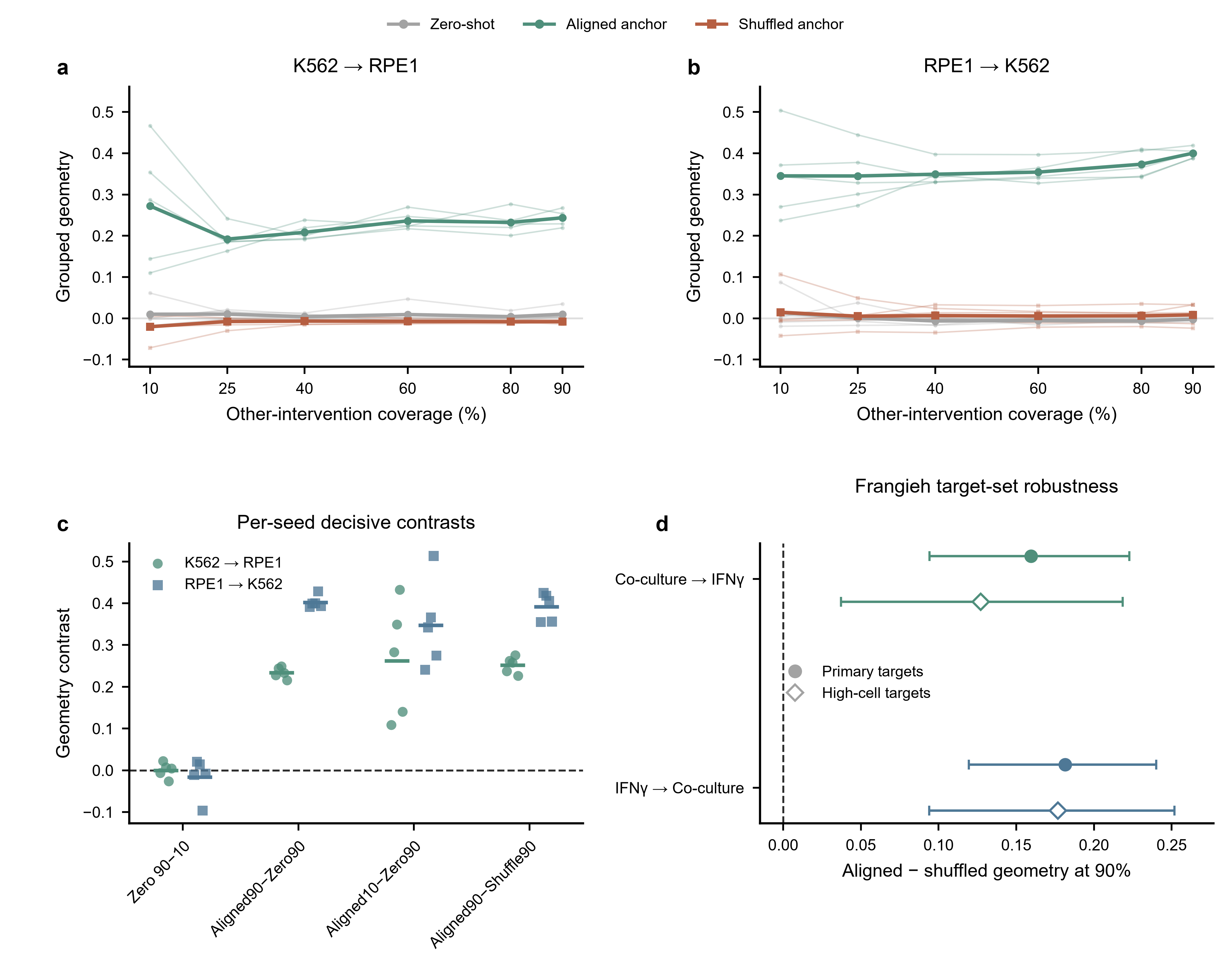

**Supplementary Figure 11 | Internal and external robustness of identity-specific empirical anchoring.** a,b, Individual frozen-seed trajectories for K562→RPE1 and RPE1→K562 across 10%, 25%, 40%, 60%, 80% and 90% other-intervention coverage. Thin lines show the five seeds and bold lines show the corresponding regime summaries for zero-shot, aligned-anchor and shuffled-anchor conditions. c, Per-seed decisive contrasts for Zero90−Zero10, Aligned90−Zero90, Aligned10−Zero90 and Aligned90−Shuffle90 in both transfer directions; short horizontal bars show the seed means and the dashed line marks zero. d, Frangieh 90% aligned-minus-shuffled contrasts for the primary target set and the predefined high-cell-count target subset in both transfer directions, with 95% confidence intervals. The figure tests robustness across internal subsampling seeds and external pseudobulk precision without repeating the aggregate coverage results already shown in Fig. 6.
